# Mutability and hypermutation antagonize immunoglobulin codon optimality

**DOI:** 10.1101/2024.03.13.584690

**Authors:** Joshua J.C. McGrath, Juyeon Park, Chloe A. Troxell, Jordan C. Chervin, Lei Li, Johnathan R. Kent, Siriruk Changrob, Yanbin Fu, Min Huang, Nai-Ying Zheng, G. Dewey Wilbanks, Sean A. Nelson, Jiayi Sun, Giorgio Inghirami, Maria Lucia L. Madariaga, George Georgiou, Patrick C. Wilson

## Abstract

The efficacy of polyclonal antibody responses is inherently linked to paratope diversity, as generated through V(D)J recombination and somatic hypermutation (SHM). These processes arose in early jawed vertebrates; however, little is known about how immunoglobulin diversity, mutability, and hypermutation have evolved in tandem with another more ubiquitous feature of protein-coding DNA – codon optimality. Here, we explore these relationships through analysis of germline *IG* genes, natural V(D)J repertoires, serum VH usage, and monoclonal antibody (mAb) expression, each through the lens of multiple optimality metrics. Strikingly, proteomic serum IgG sequencing showed that germline *IGHV* codon optimality positively correlated with VH representation after influenza vaccination, and *in vitro*, codon deoptimization of mAbs with synonymous amino acid sequences caused consistent expression loss. Germline V genes exhibit a range of codon optimality that is maintained by functionality, and inversely related to mutability. SHM caused a load-dependent deoptimization of *IGH* VDJ repertoires within human tonsils, bone marrow, and lymph nodes (including SARS-CoV-2-specific clones from mRNA vaccinees), influenza-infected mice, and zebrafish. Comparison of natural mutation profiles to true random suggests the presence of selective pressures that constrain deoptimization. These findings shed light on immunoglobulin evolution, providing unanticipated insights into the antagonistic relationship between variable region diversification, codon optimality, and antibody secretion; ultimately, the need for diversity takes precedence over that for the most efficient expression of the antibody repertoire.

## Introduction

Germline V(D)J gene diversity serves as the foundation for immunoglobulin repertoire synthesis. To support B cell receptor (BCR) generation, humans encode 46 functional immunoglobulin variable (*IGHV)* genes, 23 diversity (*IGHD)*, and 6 joining (*IGHJ)* genes as constituents for heavy chain rearrangement, as well as numerous *IGKV*, *IGKJ*, *IGLV* and *IGLJ* genes for light chain rearrangement^1^. In total, splicing of these genes through V(D)J recombination^2–5^ and junctional diversification^6,7^ can produce a theoretical ∼10^15^ unique BCR configurations^8^, facilitating enormous breadth of antigen recognition. In addition, somatic hypermutation (SHM) of V(D)J regions within activated germinal center (GC) B cells can further modify BCR variable regions^9–12^, enhancing their binding affinity^13^, flexibility^14^, adaptability^15^, and ultimate effector potential. Overall, while immunoglobulins all possess a highly conserved function in antigen recognition, variable region sequence diversity is vast and critical in maintaining human health. In balance, this diversity results in relatively few antibody-secreting cells of any specificity; thus, for effective serum protection, there also comes a requirement for extensive antibody protein secretion on a per cell basis, for which antibody mRNA must also be well adapted.

Recently, it has been established that codon usage patterns can modify the translational stability of mRNA in human cells^16–18^ following earlier studies in yeast, zebrafish, and *Xenopus*^18–20^. Specifically, when the frequency of a given codon within a transcript was correlated against the half-life of that transcript across large datasets, codons were identifiable as (de)stabilizing to varying degrees, and assigned codon stability coefficients (CSCs)^19^ equivalent to their Pearson correlation. Synonymous deoptimization of coding DNA was shown to attenuate recombinant protein expression^16^ through transcript degradation^21,22^. Ultimately, these findings demonstrate that mRNA codon optimality is an important encoded determinant of protein expression efficiency, alongside a number of other factors such as structural minimum free energy (MFE), translation initiation energy, GC%, avoidance of mRNA:ncRNA interactions, and post-transcriptional modifications (e.g. m^6^A methylation)^23–25^. To note, the concept of CSCs expands an existing definition of codon optimality, which has classically relied on analysis of contextual synonymous codon usage relative to global codon usage frequencies (CUFs) for a given species, calculated in the form of codon adaptation indices (CAIs)^26^.

Ultimately, this relationship between codon content and mRNA longevity led us to wonder whether B cell V(D)J coding regions, being diverse to an extreme, exhibit natural variability in codon optimality. Furthermore, we hypothesized that optimality may be systematically related to immunoglobulin-specific subgenomic characteristics, such as encoded mutation potential and/or post-activational mutational load. In the current study, we measured codon optimality in human germline immunoglobulin genes and natural BCR repertoires using various CSC– and CUF-based metrics. In doing so, we find that antibody V genes exhibit a range of optimality that is maintained under the selective pressures of functionality, and that both mutability and SHM antagonize immunoglobulin codon optimality. The effect of SHM was strongest in *IGH* VDJ repertoires, observable within diverse human tissues (tonsils, mediastinal & axillary lymph nodes [MLNs/ALNs], bone marrow [BM]), noted in large SARS-CoV-2-specific clonal families, and conserved in both mice and zebrafish. These findings were made across in-house and public datasets. *IGHV* optimality positively correlated with VH representation among IgG clonotypes detected in the serological repertoire of influenza vaccinees, and importantly, targeted V(D)J deoptimization of monoclonal antibodies (mAbs) caused significant expression loss. Overall, these data provide context as to how the immunoglobulin biosystem has evolved to accommodate extreme levels of sequence diversity, both germline and hypermutative, within the landscape of codon optimality.

## Results

### Immunoglobulin V(D)J genes exhibit germline variability in codon optimality

To begin, we sought to quantify baseline variability in codon optimality across human germline *IG* genes. Pre-spliced *IGHV*, native *IGHD* and *IGHJ*, and CH-S-spliced *IGHC* reference sequences were acquired from the International Immunogenetics Information System (IMGT) database^1^. Only complete, functional alleles were considered. For *IGHD* genes, all productive frames (+1/2/3) were considered to account for junctional diversification. Each sequence was segmented into individual codons, which were then assigned CSCs based on means generated from six optimality assays performed by Wu *et al*.^16^ across HEK293T, HeLa, RPE, and K562 cells (**Table S1**). CSC values ranged from 0.1304 (ATC) to –0.1485 (CAT). Subsequently, the mean over length was taken to generate CSC averages (CSC^Wu^ averages unless otherwise indicated; **Figure 1a**). The proportion of suboptimal codons (CSC < –0.01) was also calculated; these values correlated strongly (**Figure S1a**). To note, CSC^Wu^ base values also demonstrated significant positive correlations with human optimality data from two independent publications; Narula *et al.*^17^ (CSC^N^; r: 0.861) calculated as a mean of CSCs from 8 assays in HEK293T cells, and Forrest *et al.*^18^ (CSC^F^; r: 0.659) calculated using CSCs from one assay in HeLa cells (**Figure 1b**; **Table S1**). Furthermore, they aligned well with human codon usage frequency (CUF) averages ([frequency/1000 codons/codon, averaged]; r: 0.257) derived from the FDA CoCoPUTs database^27^ using RefSeq reference data (**Figure 1b**).

**Figure 1.**
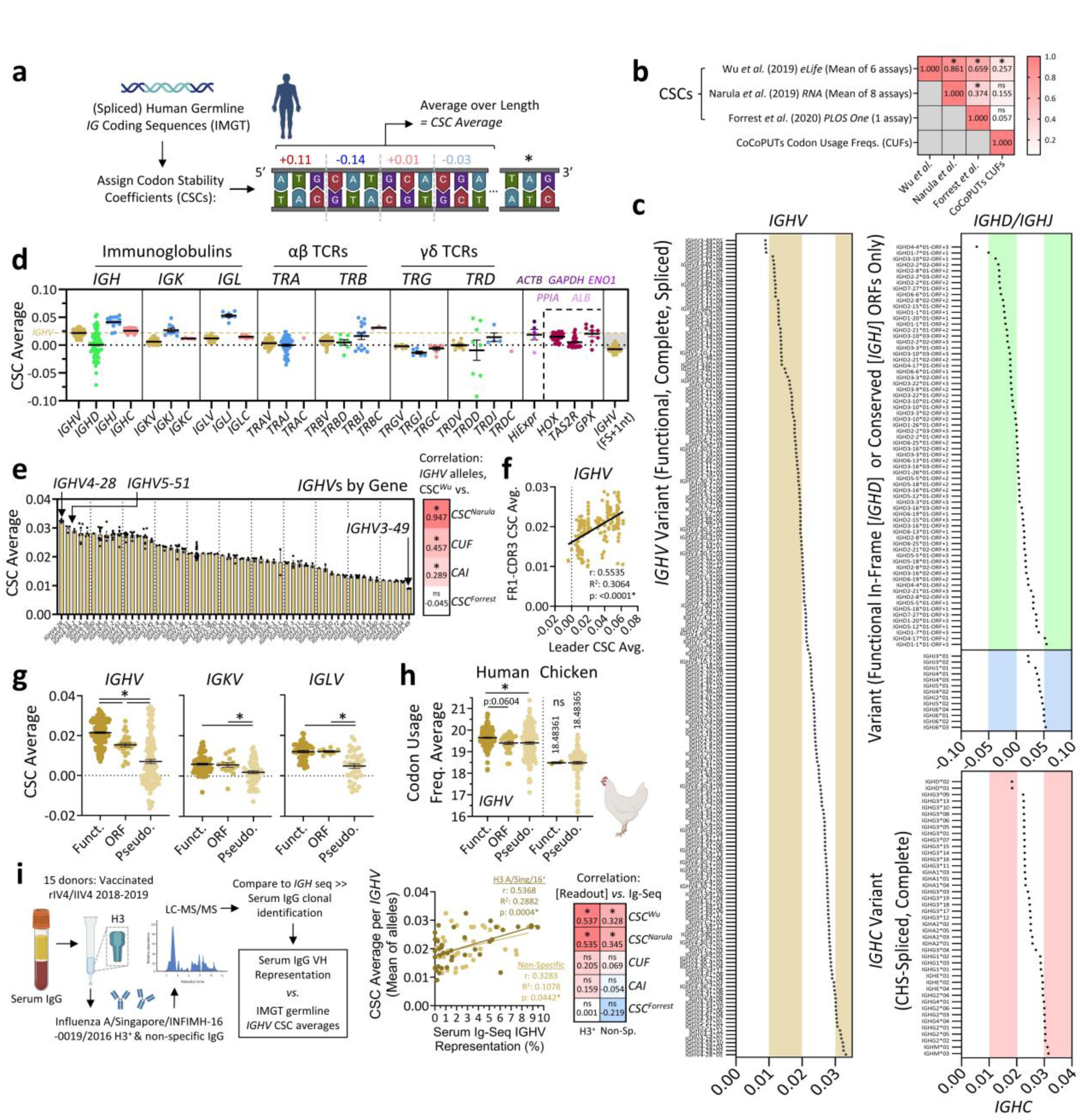
Germline immunoglobulin genes exhibit variability in codon optimality. (**a**) Human germline *IG* sequences were divided into individual codons. CSCs (based on assay means from^16–18^) or CUFs (CoCoPUTs^27^ usage frequencies per 1000 codons) were then assigned and averaged over the length. Alternatively, CAIs^26^ were calculated. **(b)** Matrix of Pearson coefficients between various CSCs and CUF scores used in this study. CAIs were not included given the discontinuous nature of the base values. **(c)** Individual CSC averages (CSC^W^, unless otherwise indicated) for functional, complete *IGH* alleles. **(d)** CSC averages for functional, complete *IG* and *TR* alleles, as well as select highly-expressed human proteins and non-immunoglobulin multigene families. **(e)** CSC averages for individual *IGHV* alleles were grouped by gene. At right, heatmap of Pearson correlation coefficients between CSC^W^ averages for each allele and other optimality metrics. **(f)** Pearson correlation of CSC values for matched *IGHV* leader sequences and FR1-CDR3 subregions. **(g)** CSC averages for functional, ORF-designated and pseudogene V gene variants were compared. **(h)** CUF averages for human and chicken *IGHV* genes. **(i)** Germline *IGHV* CSC averages were correlated against VH representation in human serum IgG from influenza-vaccinated participants. n=15 donors pooled from three groups (rIV4-, eIIV4-& ccIIV4-vaccinated). At right, heatmap of Pearson correlation coefficients between serum IGHV representation and all metrics. **Data: (d)** Mean; **(g,h)** mean ± SEM. **Statistics: (b,e,f,g[corr.])** linear regression with Pearson’s correlation; **(g,h)** one-way ANOVA with Tukey’s test. *: p<0.05. Graphics in **(a,h,i)** from Biorender.com.

For *IGHV* alleles, CSC averages ranged from 0.0089 to 0.0331, with a mean of 0.0218 (**Figure 1c/d**); *IGHD* open-reading frames (ORFs) exhibited a lower mean (0.0004) but much wider range, while *IGHJ* alleles exhibited a higher mean (0.0414) and similar range. *IGHC* gene variants demonstrated a similar mean to *IGHV*s (0.0257), and the narrowest overall range. To note, *IGHV* sequences with greater optimality tended to be slightly shorter, while *IGHJ* alleles demonstrated the opposite relationship (**Figure S1a**). *IGK* and *IGL* genes displayed similar optimality trends to *IGH*, with J regions having the highest mean optimality, C regions intermediate, and V regions the lowest (**Figure 1d**). This effect was especially pronounced for *IGLJ* genes, which were exceptionally codon-optimal compared to *IGHJ* and *IGKJ*. Importantly, all *IG* alleles (with the exception of some *IGHD* ORFs) fell within the range of codon optimality associated with a selection of mRNAs from highly-expressed proteins (actin B, GAPDH, albumin, cyclophilin A, and alpha-enolase) as well as other non-immunoglobulin multigene families (homeobox (HOX) proteins, type-2 taste receptors (TAS2Rs), and glutathione peroxidases (GPXs); **Figure 1d; Table S2**). When *IGHV* alleles were frameshifted +1bp, however, inherently disrupting the naturally-evolved ORFs, a drastic reduction in codon optimality was observed (**Figure 1d/S1b**); the same was noted for 76/77 tested *HOX*, *GPX*, *TAS2R* and highly-expressed genes, with the exception being *ALB*. In comparison to *IG* genes, the codon optimality of T cell receptor (*TR*) V(D)J alleles was generally lesser, and less dynamic, but within a similar range.

When *IGHV* alleles were grouped, we observed a high degree of consistency within genes; for instance, *IGHV4-28*, *IGHV5-51*, and *IGHV4-38-2* sequences had the highest codon optimality, while *IGHV3-49*, *IGHV3-64*, and *IGHV3-15* had the lowest (**Figure 1e**). Notably, CSC averages of individual alleles correlated well with CSC^N^ averages, CUF averages, and codon adaptation indices (CAIs)^26^ generated from usage frequencies, but not CSC^F^ averages (**Figure 1e**). Aggregate ranking using all metrics suggested that *IGHV5-51*07* was the most codon-optimal allele, and *IGHV3-35*02*, the least (**Table S3**). Within spliced *IGHV* genes, the codon optimality of leader sequences correlated positively with optimality within FR1-CDR3s (framework region 1 through complementarity-determining region 3), suggesting concurrent evolution of these subgenomic elements (**Figure 1f**).

Furthermore, we compared the optimality of functional vs. pseudogene and ORF-designated^1^ (non-functional) V gene variants. All gene groups (*IGHV*, *IGLV*, *IGKV*, *TRAV*, *TRBV*) showed a substantial decrease in codon optimality between functional variants and pseudogenes, with *IGHV* and *TRBV* alleles exhibiting the largest deficits **(Figures 1g/S1c)**. For *IGHV*s, this change was conserved regardless of which optimality metric was used (**Figures 1h/S1d**), as well as when pseudogenes were restricted to a similar length range as complete functional variants (≥341bp; **Figure S1e**). In comparison, only *IGHV* genes showed a significant reduction between functional and ORF sequences. Expanding on this data, we compared the difference between functional and pseudogene *IGHV* variants between humans and chickens (using respective CUF averages; no chicken CSC data available to date), whose pseudogenes, unlike humans, are used as functional substitutes for BCR synthesis through gene conversion^28–30^. In doing so, we observed that chicken pseudogenes exhibited no difference in mean optimality relative to functional variants, while humans did (**Figure 1h**). In aggregate, these data suggest that human V gene variants, especially those of D-containing chains (heavy BCR, β TCR), fail to maintain optimized codon repertoires in the absence of usage-associated evolutionary pressures.

We next sought to understand the *in vivo* significance of germline *IG* gene optimality. Specifically, we asked whether variation in *IGHV* optimality correlates with IgG abundance in human sera, supporting a direct impact on expression in the antibody repertoire. To test this, we compared the mean CSC average of individual *IGHV* genes to their respective abundance detected in the serum IgG repertoires of n=15 middle-aged, female subjects on day 28 following seasonal influenza vaccination. The amount of each VH type was determined by LC-MS/MS proteomics-based Ig-Seq analysis, quantifying the extracted ion chromatogram (XIC) peak area of CDRH3 peptides identified from serum clonotypes utilizing the same VH gene usage, as reported previously^31–33^. In doing so, we observed that VH types with higher codon optimality tended to show higher serum abundance among both hemagglutinin (HA)-specific (vaccine H3 strain-matched) and non-specific antibody fractions (**Figure 1i**). This relationship was maintained when CSC^N^ averages were used; positive, non-significant trends were also observed for H3-specific clonotypes using CUF averages and CAIs. In comparison, when individual H3-specific antibody clones were analyzed, we observed a weak inverse relationship between *IGH* V(D)J optimality and clonal abundance (**Figure S1f**), and no relationship for non-specific clones. Overall, these data show that differences in *IGHV* mRNA codon optimality correlate with, and suggest influence on, global serum IgG VH representation.

### Immunoglobulin V genes exhibit an inverse relationship between mutability & codon optimality

Immunoglobulin V genes are highly evolved^34–37^ to serve as a substrate for Activation-Induced Deaminase (AID)-driven hypermutation^11,38^ that is critical for variable region affinity maturation and production of high affinity protective antibodies. This occurs in part by targeting AID activity to the hypervariable regions of V genes that encode antibody CDRs, and avoiding mutation of framework regions that would be structurally detrimental^37^. Moving forward, we sought to determine whether codon optimality of germline immunoglobulin genes was affected by their predilection to act as a substrate for SHM. Only V genes were considered for this analysis given the relatively short length of D/J segments. The SHazaM package^39^ in R provides 5-mer context tendencies of somatic mutation targeting that is empirically derived from thousands of variable gene sequences^40^. Using SHazaM, mutability scores were calculated for the central nucleotides of iterative 5-mers within each sequence and the average was taken over the length (**Figure 2a**). In comparing optimality and mutability, robust inverse relationships were observed for *IGHV*, *IGKV*, and *IGLV*, indicating that the more mutable the allele, the less codon-optimal. Interestingly, similar observations were made for *TRAV* and *TRBV*; despite being non-mutative in humans, both classes of receptor may have evolved from an AID-targeted mutating precursor^41–43^. The correlation within immunoglobulin V genes was maintained when CSC^N^, CUFs, and (in the case of light chains) CAI values were instead used to quantify optimality (**Figure 2a**). In contrast, CSC^F^ averages provided an opposing, positive correlation for *IGHV*s.

**Figure 2.**
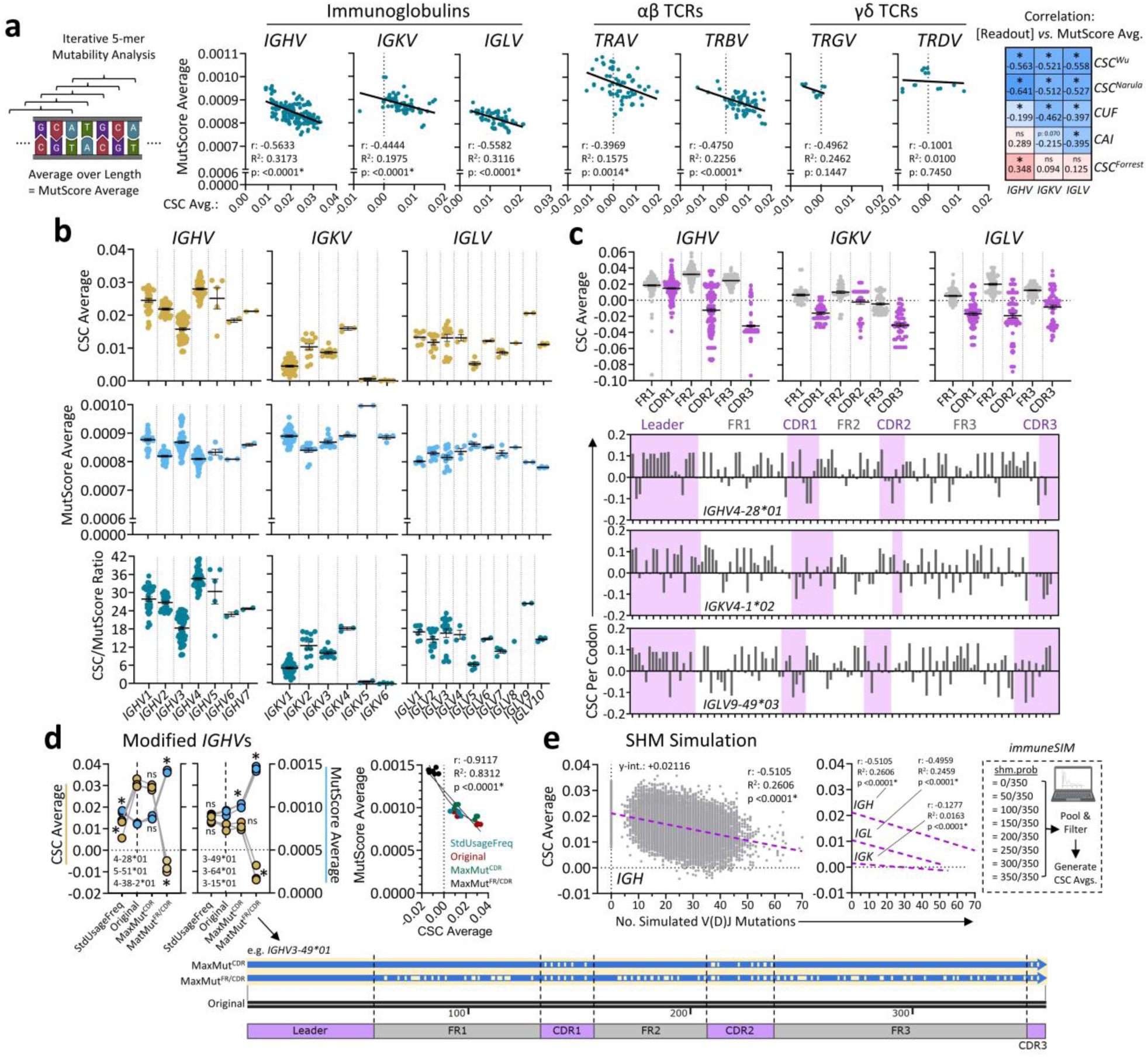
Inverse relationship between mutability and codon optimality in antibody V genes. (**a**) Mutability score (MutScore) averages were calculated for V genes using SHazaM, and correlated against CSC averages. At right, heatmap of Pearson coefficients between MutScore averages and all optimality metrics for *IGHV*, *IGKV* and *IGLV* sequences. **(b)** CSC averages, MutScore averages, and CSC/MutScore ratios were plotted for each *IG* V gene family. **(c)** Available FR and CDR regions were downloaded for each *IG* V gene sequence, and CSC values were calculated. **(d)** Maximally-mutable (CDR or FR/CDR) and standard usage frequency-aligned variants were generated for the indicated *IGHV*s. MutScore and CSC averages were then calculated and for each plotted for each matched variant. At right, correlation between CSC and MutScore average for all variants. **(e)** immuneSIM was run on reference *IGH*, *IGK* & *IGL* repertoires under varying SHM probabilities. Sequences were pooled per chain; CSCs averages were calculated and plotted against the number of mutations per sequence. **Data: (c,d)** Mean ± SEM. **Statistics: (a,d,e[corr.])** linear regression with Pearson’s correlation (non-linear regression also presented in **[d]**); **(d)** repeated measures one-way ANOVA with Geisser-Greenhouse correction and Tukey’s test. *: p<0.05. Graphics in **(a,e)** from Biorender.com.

When alleles were binned by gene family, clear tendencies were observed (**Figure 2b**). For instance, while *IGHV1*/*2* alleles showed similar trends between optimality and mutability (*IGHV2* having a lower mean optimality and mutability than *IGHV1*), *IGHV3* alleles exhibited an exceptionally low optimality and relatively high mutability, while *IGHV4* alleles exhibited the opposite. When quantified as a CSC/MutScore ratio, *IGHV1* and *IGHV2* ranked similarly, while *IGHV3* alleles had an exceptionally low ratio (more mutable), and *IGHV4* alleles an exceptionally high ratio (more codon-optimal). Overall, these data suggest that *IGHV* gene families exhibit different predilections for mutability and optimality, with *IGHV3* favouring mutation, *IGHV4* optimality, and others being moreso balanced. To note, kappa and lambda light chain genes also demonstrated variation in this ratio, with *IGKV4* and *IGLV9* representing the most optimal, and *IGKV6*/*IGLV5*, the most mutable. Broad trends in *IGHV* optimality/mutability were maintained when optimality was calculated using CSC^N^, CUF, and CAI values, but not CSC^F^ averages (**Figure S2a**).

Given the established relationship between optimality and mutability on the level of whole genes, we next asked whether CDRs, being highly mutable^34,35,37^ in determining antigen-binding affinity, exhibit differences in optimality relative to less mutable FRs. In this regard we observed that CDRs seemed to exhibit systematically lower optimalities than FRs (**Figure 2c**). The magnitude of this deficit varied; CDRH1 and CDRK2 were only marginally decreased, while the remainder were substantially lower. Similar global trends were observed for *IGHVs* when CSC^N^, and CUF were used to assess optimality, but not CAI or CSC^F^ averages; however, CDR3s demonstrated consistently reduced optimality by all metrics (**Figure S2b)**. Overall, these data suggest that mutability negatively impacts codon optimality even at the level of genomic subregions, especially within germline *IGHV* CDR3s, favouring diversity over expressibility.

We next sought to understand how *in silico* modification of immunoglobulin sequences to either increase mutability, or emulate hypermutation, affects codon optimality. First, we generated maximally-mutable, amino acid-conserved sequence variants of six germline *IGHV* sequences, three with high optimality (*IGHV4-28*01*, *IGHV5-51*01*, *IGHV4-38-2*01*) and three with low (*IGHV3-49*01*, *IGHV3-64*01*, *IGHV3-15*01;* **Figure 2d**). In one iteration, only CDRs were modified (MaxMut^CDR^); in another, both FRs/CDRs were (MaxMut^FR/CDR^). In addition, we generated variants in which codon usage was modified to reflect standard human usage frequencies in order to neutralize mutability (**Table S4**). While modifying CDRs in isolation did not significantly affect CSC averages, MaxMut^FR/CDR^ variants showed drastically reduced optimality (**Figure 2d**), supporting the natural inverse relationship within whole sequences. Furthermore, when standard codon frequencies were used to neutralize mutability, optimality and mutability scores again moved in opposition. Mutability and optimality were negatively correlated over all variants. Secondly, we ran immuneSIM^44^, an R package designed to simulate SHM on reference immunoglobulin repertoires. In modeling hypermutation of V(D)J sequences at diverse probabilities (**Table S5**), we predicted that increasing mutational load should linearly attenuate codon optimality in *IGH*, *IGL*, and to a lesser extent, *IGK* repertoires (**Figure 2e**). In aggregate, these data suggest that mutability and codon optimality are inversely related within germline immunoglobulin V genes and predict that SHM can further deoptimize recombined V(D)Js.

### Somatic hypermutation antagonizes codon optimality within natural *IGH* VDJ repertoires

To validate our *in silico* predictions, we analyzed three natural BCR datasets derived from 5’ single cell sequencing: two generated in-house (pediatric tonsil & adult mediastinal lymph node [MLN] B cells) and one public dataset from Kim *et al*.^45^ (derived from adult bone marrow [BM] plasma cells [PCs] that have been selected for antibody secretion; **Figure S3a**). Sequences were parsed for V(D)J sequence information spanning FR1-FR4 through IgBlast alignment; B cells from the pooled left & right tonsils of a female pediatric donor were analyzed first. When CSC averages were calculated, systematic differences were observed between tonsil *IGH*, *IGK* and *IGL* sequences, with *IGH* being the most codon-optimal and *IGK* the least (**Figure 3a**). Mutational differences were minimal but statistically significant; mutated *IGH* and *IGL* clones had a marginally lower optimality than unmutated clones, while mutated *IGK* clones had a marginally higher optimality. The exceptionally low optimality distribution of *IGK*s is likely attributable to the predominance of *IGKV1* variants (**Figure 2b**). To explore this further, we correlated the number of V(D)J mutations against CSC average for each sequence. As predicted by immuneSIM, we observed that *IGH* sequences with more mutations tended to have lower codon optimality. In comparison, *IGLs* demonstrated only a weak negative correlation, and *IGKs*, and a weak positive correlation (**Figure 3b**). *IGH* findings were maintained when mutational density was analyzed in place of load (**Figure S3b**), and when CSC^N^, CUF, CAI, and CSC^F^ values were used to measure optimality (**Figure 3b**). In comparison, disparate results were observed for light chains between metrics; *IGKs*, for instance, demonstrated a consistent positive relationship among CSC-based metrics (in contrast to immuneSIM predictions), but a significant negative relationship among CUF-based metrics (CUF averages, CAIs).

**Figure 3.**
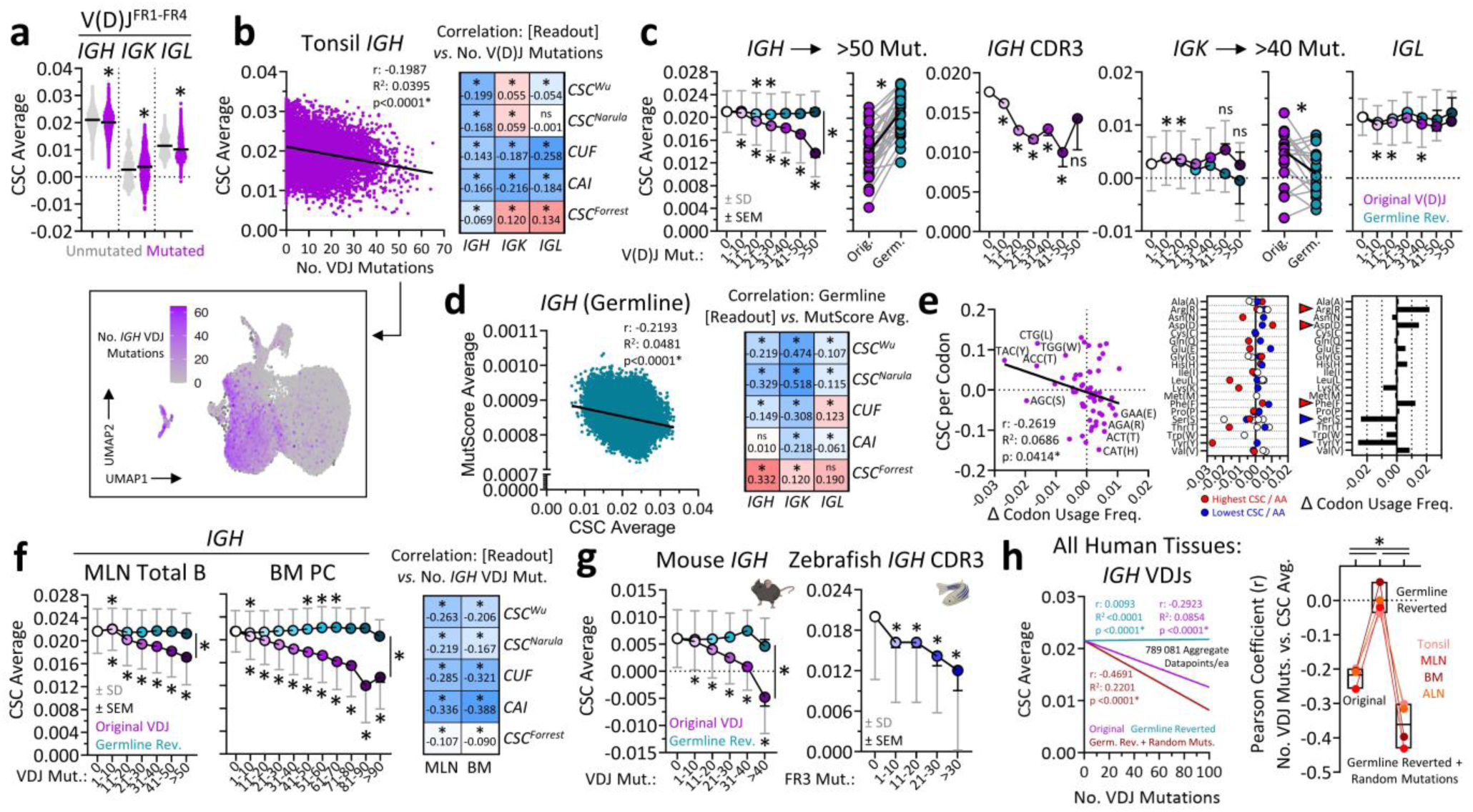
Somatic hypermutation antagonizes *IGH* VDJ codon optimality. (**a**) Total B cells from tonsils were analyzed by 5’ scRNA-seq, and CSC averages were calculated for V(D)J^FR1-FR4^ sequences. Mutated (≥1 mutation(s) per IgBlast) were compared to unmutated sequences for each chain. **(b)** Tonsil *IGH* mutational load is shown by UMAP (bottom). At middle, *IGH* CSC averages were correlated against mutational load. At right, heatmap of Pearson coefficients between mutational load and all optimality metrics for *IGH*, *IGK* and *IGL* repertoires. **(c)** Tonsil sequences (complete *IGH*, *IGK*, *IGL,* or extracted *IGH* CDR3s) were binned by mutational load; CSC averages were plotted for each sequence (purple) and corresponding germline reverted sequences (green). Additionally, matched original and germline CSC averages were plotted as paired data for highly-mutated *IGH* & *IGK* sequences. **(d)** MutScore averages were calculated for tonsil *IGH* germline sequences and correlated against CSC averages. At right, heatmap of Pearson coefficients between MutScore average and all optimality metrics for *IGH*, *IGK* and *IGL* repertoires. **(e)** Tonsil *IGH* sequences with >50 or zero mutations were pooled separately. Codons were counted for each pooled group, and usage frequencies were calculated. At left, change (Δ) in usage was correlated against the CSC per codon. At middle, change in usage for each synonymous codon was plotted. Red dots represent the most optimal codon (highest CSC), and blue dots the least. At right, synonymous codon counts were pooled, and the change in total amino acid usage frequency was plotted. **(f)** CSC averages of original and germline-reverted *IGH* sequences are plotted for MLN B cells (sequenced in-house) and BM PCs (public dataset^45^). At right, heatmap of Pearson coefficients between mutational load and all optimality metrics for MLN and BM *IGH* repertoires. **(g)** At left, pooled mouse (n=5; 34wpi H1N1 NL/09) splenocyte *IGH* repertoires (sequenced in-house) were binned by mutational load; CSC averages were plotted for original and germline variants. At right, whole zebrafish (n=1) *IGH* CDR3s (public data^48^) were binned by mutational load, and CSC averages were plotted. **(h)** *IGH* VDJ repertoires from tonsil, MLN, ALN, and BM datasets were analyzed for mutational load, and reverted to germline. Matching numbers of random mutations were then introduced into each germline sequence (n=10sequence). At left, correlations between mutational load and CSC average are presented for original, germline, and randomly-mutated pooled sequences. At right, paired Pearson coefficients are presented for each dataset/variant. **Data: (a)** Means; **(c,f,g[binned])** mean ± SEM (black) ± SD (grey); **(h)** Means, min & max. **Statistics: (a)** Unpaired t-tests; **(b,d,e,f,h[corr.])** linear regression with Pearson’s correlation; **(c,f,g[binned])** one-way ANOVA with Tukey’s test, separately for original/germline sequences. **(c[paired])** paired t-test; **(h)** repeated measures one-ANOVA with Geisser-Greenhouse and Tukey’s test. *: p<0.05. Graphics in **(g)** from Biorender.com.

When sequences were binned by mutational load, we similarly observed a progressive decrease in optimality for *IGH* VDJs and CDR3s (**Figure 3c**). Upon reversion of sequences to germline (except junctions), *IGH* optimality remained largely static over the mutational range, demonstrating that observed changes were directly due to mutations as opposed to, for instance, *IGHV* selection within highly-mutated sequences. When *IGK* and *IGL* sequences were binned, neither showed a significant difference in optimality between most mutated and unmutated, although most-mutated *IGK* sequences showed significance in reversion to germline (**Figure 3c**). To note, in support of relationships observed among V genes (**Figure 2a**), germline tonsil V(D)J sequences showed a tendency for mutability to antagonize optimality (**Figure 2d**); these effects were similarly observed in 3-4 out of 5 total metrics, with CSC^F^ analyses showing a consistent disparate effect (**Figure 2d**).

We also considered other correlates of selection, including isotype switching and phenotype. Compared to IgD^+^ and IgM^+^ cells, *IGH* VDJ sequences from IgA1^+^, IgA2^+^, IgG1^+^ and IgG2^+^ B cells displayed marginally but significantly lower optimality (**Figure S3c**). Similarly, specific cell phenotypes were also attenuated by a similar magnitude, such as *FCRL4/5*^lo^ class-switched and non-class-switched memory B cells ([n]cs-MBCs), *FCRL4/5*^hi^ csMBCs^46^, and plasmablasts. However, these suppressive effects appeared to be dependent on co-varying mutational load; analysis of unmutated sequences reported no difference between most isotypes (except IgG1, which was marginally increased vs. IgD) or clusters (except 5, 7, and 8 which were minorly increased relative to cluster 0; **Figure S3c**). There was no significant enrichment of highly optimal *IGHVs* among naïve B cells (**Figure S3d**).

The negative correlation between mutational load and *IGH* optimality suggested that specific codons were being systematically enriched during SHM. In analyzing codon counts for highly-mutated (>50 mutations) compared to unmutated VDJs, it was observed that a select few codons were greatly depleted (>–0.01 change in codon usage frequency; **Figure S3e; Table S6**). These included TAC (Tyr), AGC (Ser), ACC (Thr), CTG (Leu), and AAG (Lys). Reciprocally, only one codon, GAC (Asp), was enriched to this level (>+0.01 change), with GAA (Glu), ACA (Thr), TTT (Phe) and AGG (Arg) coming close. As expected, the majority of codons contained within the classical AID hotspot^34,35,37^ motif DGYW/WRCH^47^ (AGT/G/CT/AT and reverse complement AT/GA/C/TAC, respectively) were depleted; three were strongly depleted (>–0.01 change; TAC, AGC, ACC), six were moderately depleted (<–0.01 change; AAC, AGT, GGT, GCT, GCA, GTA), two were neutral (TGT, TGC), while five were moderately increased (<+0.01 change; ACA, ACT, GTT, GCC, GGC). Notably, when change in frequency was correlated against the individual CSC per codon, it was apparent that lowly-optimal codons were enriched, and highly-optimal codons depleted during SHM (**Figure 3e**). Furthermore, when we plotted change in frequency by amino acid, we observed that for 12/18 multi-codon amino acids, the most optimal codon was the most depleted (or least enriched). Similarly, for 8/18, the least optimal codon was the most enriched. Overall, in terms of amino acids, Arg, Asp and Phe were consistently increased in highly-mutated clones, while Ser, Tyr, and Lys were decreased. These findings show that functional mutation targeting in conjunction with selection during affinity maturation favors diversity over expression optimization.

We next analyzed two additional human datasets: pooled MLN B cells from the lungs of an adult organ donor, and a public dataset^45^ comprising BM PCs from 11 COVID mRNA vaccine recipients (**Figure S3a)**. In line with tonsils, both repertoires showed that higher *IGH* VDJ mutational load was associated with reduced codon optimality (**Figure 3f/S3f**). Correlations were highly similar for all three datasets, demonstrating a clear antagonistic relationship between hypermutation and optimality that was supported by all optimality metrics (**Figure 3f**). Furthermore, changes in both codon and amino acid frequencies correlated robustly under multilinear regression (**Figure S3g; Table S6**). Finally, to complement our analyses of human repertoires, we assessed mutational codon deoptimization in two animal models: mice and zebrafish (*D. rerio*). For mice, we performed 5’ scRNA-seq on splenic B cells collected at 34 weeks-post infection with influenza H1N1 A/Netherlands/602/2009, and calculated optimality using mouse CSC values from Forrest *et al*.^18^ (**Table S1**). For zebrafish, we analyzed a public dataset from Weinstein *et al*.^48^ featuring short-read heavy chain repertoires from whole fish, and calculated optimality using zebrafish CSC values from Bazzini *et al*.^20^ (**Table S1**). Because zebrafish reads were partial, we generated CSC averages for CDR3s, and used IMGT/HighV-QUEST to infer mutational load within matched FR3s. For both species, we found significant negative relationships between *IGH* mutation and codon optimality (**Figure 3g**), in line with our findings in human lymphoid organs.

In trying to better understand why some sequences deoptimized to different degrees in response to the same number of mutations, we hypothesized that the magnitude of change in optimality related to the germline optimality of the *IGHV* used. To test this, we filtered BM PCs (the largest dataset) to assess cells with a fixed mutation count (40; selected to balance sequence number, diversity and mutational load). In doing so, we found a significant positive correlation suggesting that more initially optimal *IGHVs* are more prone to deoptimize than less optimal ones, when mutated to the same degree (**Figure S3h**). Tonsil and MLN B cells demonstrated similar trends, although non-significantly due to smaller dataset sizes and reduced diversity of clones with a fixed mutation count. Overall, however, these data demonstrate that SHM systematically antagonizes the codon optimality of *IGH* VDJ regions across multiple species, with greater effect in sequences with initially-optimal *IGHV* genes.

### Natural restriction of mutational codon deoptimization

To test how much natural SHM deoptimizes VDJs compared to random mutation, each human *IGH* dataset (tonsils, MLNs (in-house), BM^45^, as well as axillary LNs^45^ [ALNs; see **Figure 4**]) was analyzed for mutational load and reverted back to germline. Subsequently, the same number of mutations were introduced back into respective germline sequences in a random (computationally pseudorandom) fashion, iterated ten times per sequence. Ultimately, in both aggregate data (left; **Figure 3h**) and individual tissues (right), randomly mutated repertoires exhibited greater deoptimization than naturally mutated repertoires. This suggests that SHM is either targeted in a manner, and/or mutated B cells selected in a manner that minimizes codon deoptimization in *IGH* VDJ regions.

**Figure 4.**
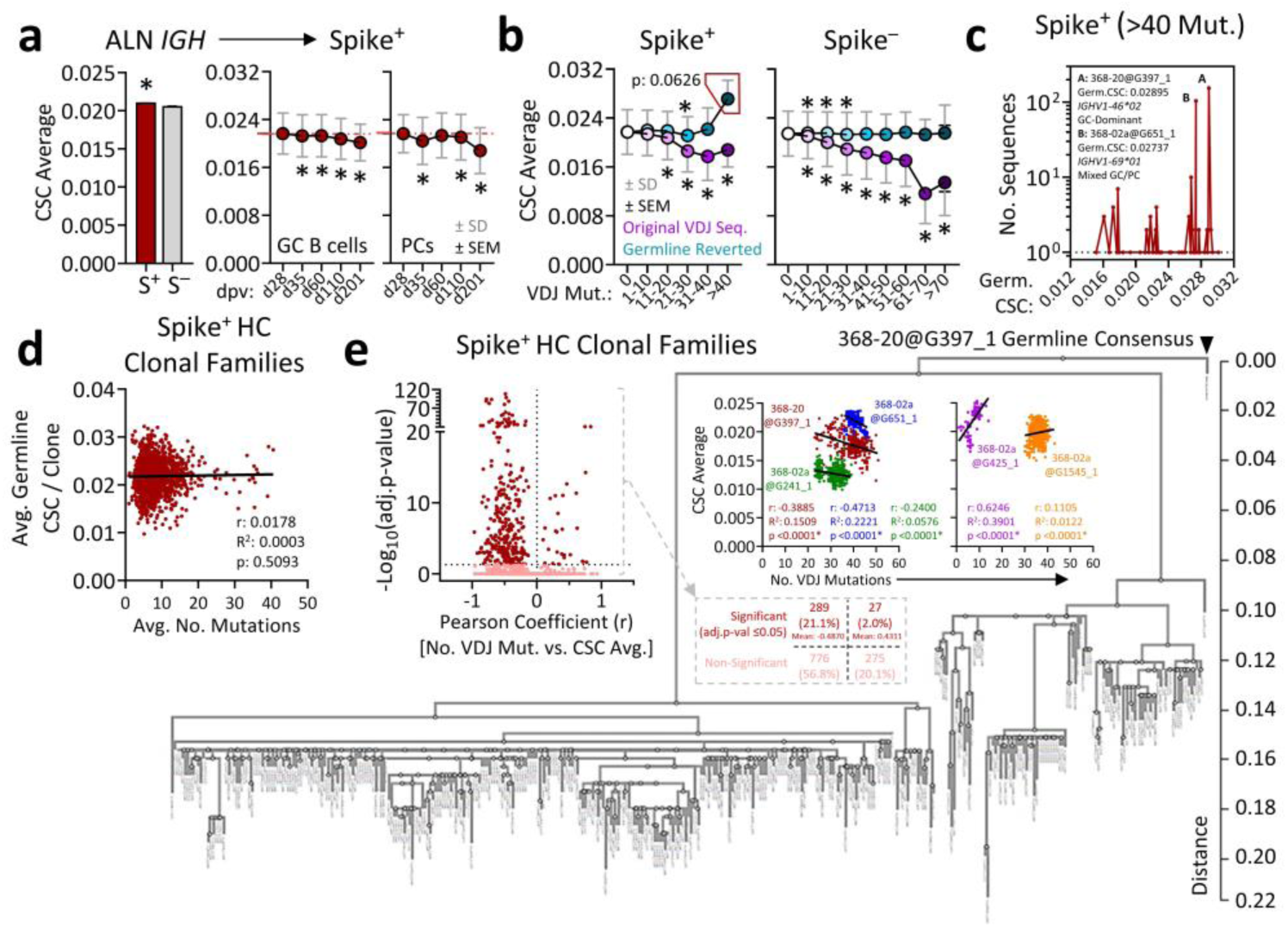
Codon deoptimization within large SARS-CoV-2-specific clonal families. (**a**) ALN B cells (public dataset^45^) were split into SARS-CoV-2 spike^+/−^ fractions as defined by Kim *et al.*^45^. At left, *IGH* CSC averages for spike^+^ B cells are compared to spike^−^ B cells. At right, CSC averages are presented at various timepoints for spike^+^ GC B cells and PCs. **(b)** Spike^+/−^ *IGH* clones were binned by mutational load; CSC averages were plotted for each sequence (purple) and corresponding germline reverted sequences (green). The red box highlights a region of interest, assessed further in (c). **(c)** ALN spike^+^ clonal families with >40 mutations were reverted to germline; subsequently, the number of *IGH* sequences with any given germline optimality score was plotted. Two highly-expanded clones are indicated. **(d)** The average germline CSC average per spike^+^ heavy chain clonal family was correlated against the average number of *IGH* mutations per clonal family. **(e)** For each spike^+^ *IGH* clonal family, VDJ mutational load was correlated against CSC average. At left, volcano plot showing families with significant negative or positive Pearson coefficients. Proportional values for this plot are presented at the end of the grey arrow. At right, representative families with negative or positive Pearson coefficients. The surrounding phylogeny is for clonal family 368-20@G397_1 (401 members). **Data: (a[bar])** Mean ± SEM; **(a[timecourse],b)** mean ± SEM (black) ± SD (grey). **Statistics: (a[bar])** Unpaired t-test; **(a[timecourse],b)** repeated measures one-ANOVA with Geisser-Greenhouse and Tukey’s test (independently for original and germline in [b]); **(d,e)** linear regression with Pearson’s correlation (and Bonferroni correction in [e]). *: p<0.05.

### Codon deoptimization in SARS-CoV-2-specific clonal families

Next, we sought to characterize changes in codon optimality following immunization. To do this, we analyzed an additional portion of the public dataset by Kim *et al*.^45^ consisting of B cells sorted from axillary lymph node (ALN) fine-need aspirates acquired at various timepoint following SARS-CoV-2 mRNA vaccination (**Figure S4a**). In aggregate, sequences previously identified^45^ as belonging to spike-reactive clonal families (S^+^) displayed a significant but negligible increase in codon optimality compared to non-reactive (S^−^) (**Figure 4a**). Over a timeframe at which mutations were shown to accumulate after vaccination^45^, there was a slight decrease in optimality for spike^+^ sequences from both germinal center (GC) B cells and plasma cells (PC; **Figure 4a**). When spike^+^ and spike^−^ sequences were compared, both displayed similar negative correlations between optimality and mutational load (**Figure S4b**), as well as decreases in binned optimality with respect to germline (**Figure 4b**).

Interestingly, we observed that within the spike^+^ group, cells which had the highest number of mutations (>40) displayed an elevated germline CSC average compared to lesser-mutated sequences (**Figure 4b**). This was not observed for spike^−^ sequences, leading us to ask whether exceptionally optimal spike-reactive clones were preferentially selected following vaccination. For sequences with >40 mutations, this difference was observed in both GC B cells and PCs (**Figure S4c**). In plotting the number of sequences with given germline CSC averages, we observed that the elevated mean (**Figure 4b**) was largely attributable to an expansion of two, relatively optimal spike^+^ heavy chain clonal families, 368-20@G397_1 & 368-02a@G651_1^45^ (**Figure 4c**). Furthermore, upon calculating mean germline optimality scores for each available spike^+^ clonal family, irrespective of mutational load, we observed no correlation with maximum/average number of mutations per family or number of clonal members (**Figure 4d/S4d**). Instead, we found that the largest clone 368-20@G397_1 contracted over time (**Figure S4e**). Overall, these data suggest that our observation of elevated germline mean optimality in highly mutated sequences (**Figure 4e**) was attributable to a transient dominance of two heavy chain clones, as opposed to a generalizable advantage for germline-optimal clones within GCs.

Despite this, these large spike^+^ heavy chain clonal families provided further opportunity to corroborate SHM-related findings. We found that sequences from both 368-20@G397_1 and 368-02a@G651_1 demonstrated significant antagonism between hypermutation and optimality (**Figure 4e**), supporting observations made at the level of whole repertoires (**Figure 3**). Furthermore, when phylogenetic relationships were inferred for 368-20@G397_1 (401 sequences), we observed a similarly robust negative correlation between optimality and phylogenetic distance to germline consensus (**Figure S4f**). While the majority of clonal families (76.9%) demonstrated no significant relationship between mutational load and optimality, those that did tended to demonstrate a decrease (21.1%) as opposed to increase (2.0%) (**Figure 4e**). In aggregate, although antigen exposure does not appear to select for exceptionally codon-optimal naïve B cell clones, hypermutation and intraclonal evolution can reduce the optimality of VDJ regions within large heavy chain clonal families.

### Relationship between V(D)J codon optimality and mAb expression efficiency *in vitro*

In preparation for mAb expression studies, we considered the fact that V(D)J regions comprise only part of BCR ORFs (**Figure 5a**), asking whether variability in V(D)J codon optimality is buffered by constant regions *in silico*. Using heavy and light chain repertoires from tonsil B cells, we compared the CSC averages of V(D)J^FR1-FR4^ sequences to the same sequences spliced with V-matched leader sequences (“LV(D)Js”) ± relevant constant regions (‘LV(D)JCs”; **Figure 5b**). We found that addition of heavy chain C regions reduced overall variability in CSC average by 69.6–75.6%, and light chain C regions by ∼45%; these magnitudes correlated strongly with the length of the C regions (**Figure 5c**). These data suggest that constant regions can partially buffer variability in V(D)J optimality.

**Figure 5.**
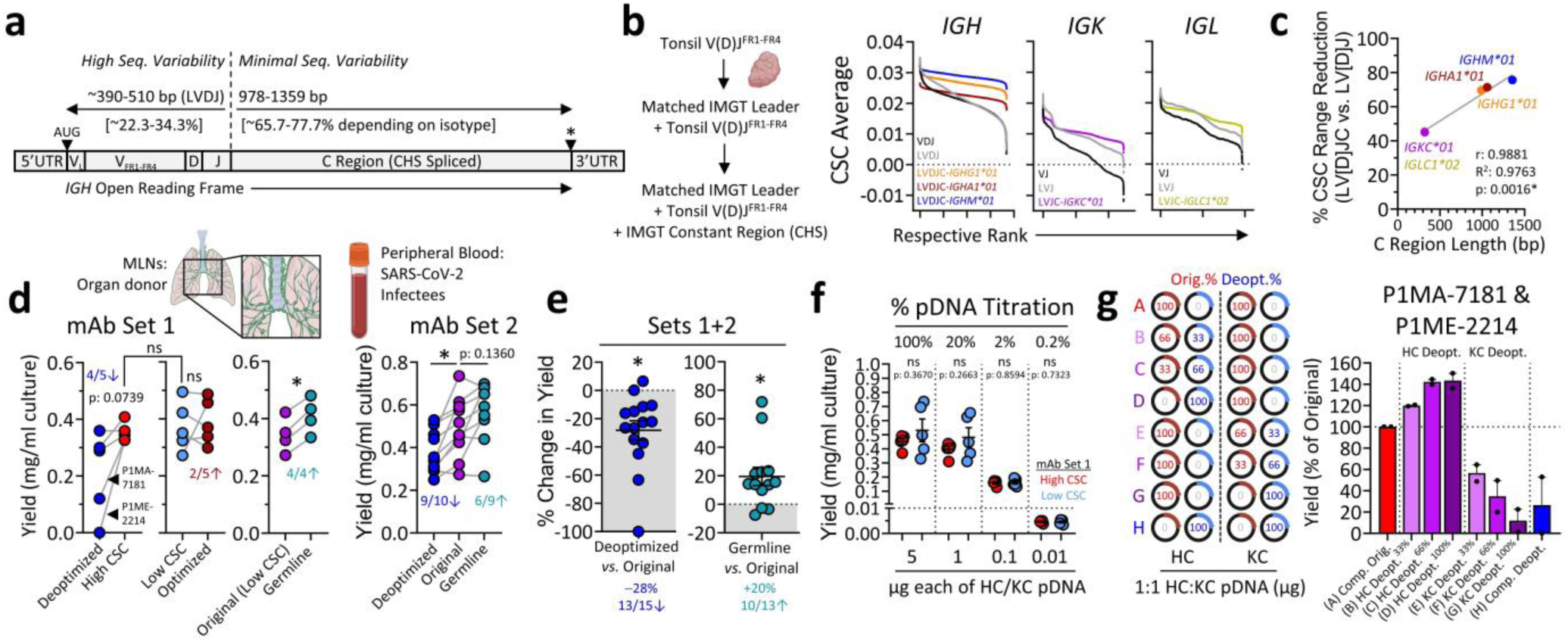
Relationship between V(D)J codon optimality and mAb expression efficiency. (**a**) Schematic demonstrating the approx. contribution of *IGH* VDJ and constant regions to heavy chain mRNA ORF length. **(b)** CSC averages for tonsil V(D)J^FR1-FR4^ sequences were ranked & plotted, before and after splicing with V-matched leader sequences ± indicated constant regions (IMGT). Each dataset was ranked independently for visual clarity. **(c)** Percent change in CSC range (max–min) elicited by constant region splicing was plotted against corresponding constant region length. **(d)** mAbs from two sets ([1]; MLN-derived, high/low natural CSC, matched (de)optimized & germline variants, and; [2] PBMC-derived, matched original, germline & deoptimized) were expressed in Expi293F cells; yield in supernatants was quantified by total IgG ELISA. Data are from one transfection per experiment. **(e)** Percent change in expression for deoptimized and germline variants relative to matched original clones for pooled sets 1 & 2. **(f)** For select mAbs (natural high/low CSC, set 1), pDNA transfection mass was titrated. **(g)** Deoptimized plasmids were titrated up for heavy & light chains independently. **Data: (e,f,g)** mean ± SEM. **Statistics: (c)** linear regression with Pearson’s correlation; **(d)** t-tests, paired where grey lines link matched clones, unpaired where comparing high/low CSC groups (bright red/blue). **(e)** one-sample t-test against theoretical mean = 0. *: p<0.05. Graphics in **(b,d)** from Biorender.com.

To test expression *in vitro*, we selected a set of mAbs from our human MLN repertoire (set 1; **Table S7**) consisting of five clones with high relative optimality across multiple metrics, and five with low (**Figure S5a/b**). Each clone had the same light chain (kappa), unique *IGHV/IGKV* pairings (**Figure S5a**), and V(D)Js of similar respective lengths (**Figure S5c**). Mutational load was unrestricted; as a result, 4/5 low optimality clones were mutated to some degree, *versus* 0/5 highly-optimal (**Figure S5d**). To note, it is difficult to compare expression levels due to codon optimality *in vitro* for unique mAbs from different B cell clonotypes because natural codon effects are likely subtle, and the translated peptide sequences can also drastically affect expression levels and would have selective influences that balance translation levels (data not shown). Differences in amino acid sequence adds additional variables such as peptide processing and post-translational modifications which can affect intracellular trafficking/secretion efficiency^49^, unspecific uptake/degradation sensitivity^50^, and stability/aggregation tendencies^51^. However, intentionally (de)optimizing codons used while maintaining the identical amino acid sequence for single B cell clones controls for this variable. Thus, we also generated additional mAb variants in which we: a) deoptimized the high CSC clones (**Figure S5e**); b) optimized the low CSC clones, and c) germline reverted the mutated clones. All V(D)Js were cloned into AbVec vectors^52^ with a uniform leader sequence to minimize differences in intracellular trafficking. Following transfection in Expi293F cells, we observed no significant difference in expression between our highly– and lowly-optimal natural mAbs (**Figure 5d**). This was maintained when uniform leader sequences were swapped for individualized, V-matched leaders (**Figure S5f**), and when pDNA transfection mass was titrated down (**Figure 5f**). V(D)J optimization did not increase expression yield of lowly-optimal clones. Importantly, however, a strong trend (p: 0.0739) towards reduced expression was observed for maximally-deoptimized synonymous clones; 4/5 deoptimized mAbs showed some expression loss, with two (P1MA-7181 & P1ME-2214) exhibiting drastically lower yield (63% and 100% respectively; **Figure 5d**). Light chain deoptimization appeared to play the dominant role in expression loss for these clones (**Figure 5g**). In comparison, germline reverted mAbs demonstrated a consistent and significant increase in expression relative to mutated parent mAbs (**Figure 5d**).

To validate these findings, we expressed an additional set of original, deoptimized and germline mAb variants (set 2) comprising ten clones identified among peripheral blood B cells from SARS-CoV-2 infected & convalescent individuals (**Table S7**). Parent clones were unrestricted with respect to light chain, *IGHV/IGKV* pairing, and V(D)J length, but all were mutated. Similar to MLN mAbs, deoptimization caused significant expression loss compared to original mAbs, while germline reversion demonstrated a trend towards increased yield (**Figure 5d**). When mAb sets were internally normalized by percent change in expression compared to original, and pooled, deoptimization resulted in diminished yield for 13/15 clones at an average of –28%, while germline reversion increased yield in 10/13 clones by an average of +20% (**Figure 5e**). Thus, in support of serum IgG VH sequencing data (**Figure 1i**), and despite the buffering capacity of constant regions, suboptimal codon usage likely affects overall antibody repertoire expression tendencies in a detrimental manner.

## Discussion

Affinity maturation^12,13^ is the elegant Darwinian process by which antigen-specific B cells adapt to cognate antigens, enhancing immunoglobulin effector capacity and honing the immune system against future threats. Somatic hypermutation (SHM)^9^ is the driving force behind this process; by mutating codons within V(D)J DNA, variable region amino acids are modified to affect paratope conformation and antibody specificity. SHM is in turn facilitated by the enzymatic activity of activation-induced cytidine deaminase (AID)^11,38^, which introduces C-to-T and G-to-A mutations within DNA “hotspot” motifs^34,35,37,47^ that have evolved within immunoglobulin genes. Our group previously reported that C/G nucleotide localization within *IGHV*s evolved in a manner that causes most primary C-to-T and G-to-A mutations elicited during SHM to be synonymous, preventing AID from introducing severe bias in amino acid shift, and A/T localization in a manner that targets polymerase-eta errors in the repair phase of SHM to CDRs^36^. Thus, most codons utilized in variable genes are affected by evolutionary pressure to support antibody diversification and affinity maturation. While it is clear that such specialized subgenomic architecture is necessary to facilitate effective SHM, this raises an interesting question: how have mutability and somatic hypermutation evolved with immunoglobulin repertoires alongside another omnipresent feature of antibody coding regions – codon optimality? As antibody secretion is critical to effective immunity, amounting to several grams of antibody protein per day in a healthy adult^53^, how is the need for optimal protein production equilibrated with that for adaptability and diversity of antibody specificities? We address this knowledge gap in the current study through analysis of human germline immunoglobulin genes and natural V(D)J repertoires.

To date, quantitative assessment of codon optimality has largely relied on analysis of codon adaptation indices (CAIs)^26^. CAIs are calculated using a reference set of usage frequencies (CUFs; usage/1000 codons; here, from the FDA CoCoPUTs^27^ database compiled from all RefSeq-indexed human data), and represent the extent to which a sequence contains infrequently-used codons. By nature, CAIs are normalized by amino acid content, being a ratio of the geometric mean of relative synonymous codon usage (RSCU) values for a given sequence relative to the geometric mean of the theoretical maximum RSCU for the same amino acids. As such, this metric represents how *synonymously* optimized a sequence is (i.e., optimized relative to its ideal codon content). While informative, CAIs inherently do not factor in differences in optimality that accompany differences in amino acid sequence. With this in mind we developed a complementary metric, the CUF average, which represents the mean of usage frequencies over the length of a sequence, does not normalize by amino acid content, and provides a readout that is more continuous between disparate ORFs. Overall, CAIs and CUF average results correlate well (e.g. for *IGHV* allele data [**Figure 1e**], r: 0.7411, p<0.0001) but represent subtly different quantitations of optimality – as defined by usage frequency.

Beyond usage frequency, however, we also felt the strong need to consider a more recently emerged definition of codon optimality – the relationship between rate of codon occurrence, experimental transcript longevity, and protein expression^16–20^. By taking the mean of codon stability coefficients (CSCs) assigned to codons over the length of a sequence, CSC averages described herein estimate the net relationship between codon content and immunoglobulin mRNA longevity. This is in comparison to CUF-based metrics (CAIs, CUF averages), which are predicated on the idea that mRNAs whose codons align well with species-specific global usage patterns are best adapted. Overall, because of the experimental data supporting CSC derivation, and the functional definition of optimality that this provides, we selected CSC averages as the primary readout in this study. Specifically, the mean of CSCs generated by Wu *et al*.^16^ were used given the robust diversity of this analysis (six results per codon, across three assay types [endogenous transcriptome decay, ORFome decay, SLAM-seq] and four human cell types [HEK293T, HeLa, K562, RPE]).

The seven *IGHV* families, defined by phylogenetic similarity^54^, displayed quite variable degrees of codon optimality despite occurring within a range defined by highly-expressed genes and other non-*IG* multigene families. Strikingly, however, we show by using Ig-Seq serum proteomic analysis that this variability is reflected in the serum abundance of IgG VH types among influenza vaccinees. This was evident among purified influenza HA-specific clones, as well as non-specific IgG. While *IGHV* optimality may co-vary with other independent factors, such as exposure history through infection/vaccination or naïve precursor frequency, these data suggest that serum IgG expression levels may be subtly influenced by differences in V(D)J codon optimality.

Mammalian pseudogenes are known to accumulate mutations and undergo genetic drift at a greater rate than functional homologues, as a result of bypassing selective pressures associated with functionality^55–57^. In comparison, chicken *IGHV* pseudogenes retain functionality in serving as donor sequences to diversify V(D)J regions during B cell development^28–30^. In line with these ideas, we observed that codon optimality was significantly reduced in human, but not chicken, pseudogenes relative to functional variants for each species. These data strongly support the notion that codon optimality is maintained in human V genes as a result of functionality-associated selective pressures.

The natural inverse relationship observed between optimality and mutability in germline V genes was of significant interest, especially given that artificial maximization and neutralization of mutability also led to reciprocal changes in optimality. Together, these findings suggest that the evolutionary accumulation of AID hotspot motifs^34,35,37,47^, or specific hotspot variants^40^, biases codon usage towards less optimal configurations. Alternatively, the accumulation of mutational “coldspots” may bias towards more optimal configurations; both factors might contribute. This finding furthermore begs the question of whether *IGHVs* evolved an elevated mean optimality to compensate for high rates of hypermutation within heavy chains (*versus* light chains)^58^. Relatedly, notable among *IGHV* families was *IGHV3*; while demonstrating a similarly high mutability to *IGHV1* genes, *IGHV3* genes had markedly lower mean codon optimality. This effect appeared to be bimodal, with especially pronounced suboptimality in two clades: one *IGHV3*-outgroup containing *IGHV3-15, –49, –72* and *-73*, and one ingroup containing *IGHV3-64*, *-64D*, *-13*, *-9, –20, –43* and *-43D* (per IMGT^1^ phylogenies). *IGHV3* genes represent the largest *IGHV* family, are the primary contributors to VH structural diversity, and are evolutionarily distinct from all other *IGHVs*, being the only members of phylogenetic clan III^59^. Thus, this data suggests that in the case of *IGHV3s*, both mutability and extensive structural diversification may have contributed to codon deoptimization. Phylogenetic analysis of *IGHV*s from multiple extant and extinct species is warranted to further understand the complex relationships between gene diversification, mutation potential, and codon optimality during *IGHV* evolution.

Upon next considering the effect of natural SHM on codon optimality, we observed a striking negative relationship within *IGH* repertoires that spanned all metrics and human tissues analyzed. In each case, increasing mutational load promoted codon deoptimization that was fully rescuable upon reversion to germline. This effect was conserved in mice, as well as the more phylogenetically distant ectotherm, zebrafish. Although consistently antagonistic, this process appeared to be at least partially constrained by specific SHM targeting and/or selection of mutated B cells, given that naturally-mutated VDJ repertoires deoptimized significantly less than randomly-mutated equivalents. In comparison, correlations within light chain sequences were more variable; CSC-based metrics tended to demonstrate mild positive relationships with mutational load, while CUF-based metrics were relatively negative. These dichotomous results (and other cases where CSC and CUF-based metrics disagree) may result from mutated light chain repertoires being enriched for codons with low usage frequency but neutral/high CSC^W/N^/CSC^F^ optimality, respectively (**Figure S6**). The clear mutational effect in *IGH* repertoires was plausibly due (at least in part) to the exceptionally high germline optimality of *IGHV* genes, being the bulk contributors to VDJ length; mean *IGHV* optimality was notably on par with many highly-expressed genes, and greater than all other *IG*/*TR* V genes and multigene families assessed. Thus, in the case of *IGHVs* it might be hypothesized that mutational perturbation could incrementally neutralize exceptional optimality. This idea is supported by the observation that sequences with highly-optimal germline *IGHVs* deoptimize to a greater degree than those with less optimal *IGHVs* under the same mutational load (**Figure S3h**). In aggregate, these data show a clear deleterious effect of SHM on codon optimality within *IGH* sequences, with more unclear effect on light chains.

To test this idea, we expressed recombinant mAbs from human repertoires, as well as synonymous (de)optimized and germline variants, under controlled conditions. To note, is difficult to demonstrate and interpret differences in expression levels between mAbs with both varied codon optimality and unique peptide sequences, the latter of which may drastically affect translation, stability, and/or secretion efficiency, and would also be selected by evolution. However, deoptimization of codons from antibody V(D)Js with identical amino acids clearly and consistently reduced expression levels, and reversion to germline increased yield in most tested clones. These results are in line with serum IgG proteomic sequencing data, and demonstrate that codon optimality may influence expression of serum antibody through *IGHV* usage. Furthermore, they support the idea that SHM can subtly compromise antibody expression efficiency. Although mechanistic inferences related to codon deoptimization are complicated by amino acid reversion in the case of germline clones, the improved yield of these mAbs demonstrates that clonal mutation can modulate expression through either mRNA– or protein-based effects. Moving forward, it may be of interest to assess the effect of V(D)J optimality within scFvs, scFabs, camelid V_H_Hs and other engineered antibody fragments^60,61^; these constructs may be more susceptible to differences in codon optimality given that V(D)J content comprises a comparatively greater total portion of their ORFs.

Somatic mutations are essential to facilitate affinity maturation and enhance antibody functionality. To this end, a reasonable null hypothesis might be that evolved hypermutative systems should not operate at the extreme expense of antibody yield. In partial contrast to this idea, we observe that *IGHV* optimality correlates with serum VH usage, and that mutability and hypermutation antagonize codon optimality within *IGH* repertoires. However, these effects appear to be partially constrained by selective pressures, as well as the buffering capacity provided by constant regions. To note, it has yet to be determined if this process occurs in the absence of, or despite, risk associated with extreme deoptimization during heavy mutation – it is possible that survivorship bias precludes detection of highly mutated clones that are selected against *in vivo* due to BCR expression loss. While we did not observe any exceptionally negative optimality outliers among apparent GC B cells (tonsil clusters 6/9/13; Figure S3e), any such highly-mutated clones are likely rare; B cells with exceptional loss of expression due to codon deoptimization may be at a selective disadvantage and therefore not detectable. Further investigation is required to explore this possibility.

To note, other recent work^62^ assessing codon usage within immunoglobulin genes showed that variable region repertoires from BMPCs and MBCs^45^ were enriched for codons dependent on flexible inosine (I)34-modified tRNAs^62^. Furthermore, they showed that antibody-secreting cells had high levels of I34-modified tRNAs, and that hyperutilization of I34-dependent codons enhanced selection for transgenic BCRs. In our analysis of the same BMPC dataset^45^, six out of eight I34-dependent codons were enriched (ATC, CCC, CGC, CTC, GCC, GTC) in highly-mutated sequences, while two were depleted (ACC, TCC; **Table S6**), supporting this^62^ observation. In comparison, this was reduced to five enriched I34-dependent codons for MLNs, and four enriched for tonsils (**Table S6; Figure S3e**). These data provide important context for our study; while we have shown that SHM systematically deoptimizes *IGH* variable regions in terms of usage frequency-and CSC-based indices, to the potential detriment of antibody expression, simultaneous enrichment for I34-dependent codons may concurrently enhance expression to counterbalance this effect in MBCs and PCs.

In summary, our work provides insight into the evolutionary relationship between mutability, SHM, and codon optimality within human immunoglobulin genes. V genes exhibited variability in optimality, but were spread within a range similar to other highly-expressed proteins and multigene families. Comparison of pseudogenes between humans and chickens showed that human *IGHV* genes maintain optimality under functionality-associated selective pressures. Furthermore, genic optimality scores correlated with VH representation in total and HA-specific IgG from influenza vaccinees, suggesting that codon use may subtly influence IgG expression *in vivo*. V gene mutability was inversely related to codon optimality, and certain gene families demonstrated biases toward one factor (e.g. *IGHV3*, mutability) or the other (e.g. *IGHV4*, optimality); artificial manipulation of mutability supported this relationship. Through analysis of multiple human datasets, we found that SHM antagonizes *IGH* codon optimality in a load-dependent manner, but that mutational targeting and/or B cell selection have evolved to minimize deoptimization relative to true random. *IGH* deoptimization was conserved in both mice and zebrafish, one of the most divergent species which retains all key features of human B cell development^48^. Although no differences were observed in the expression of natural mAbs with high/low V(D)J optimality *in vitro*, antibodies with distinct amino acid sequences will undergo selection based on additional factors at the protein level (e.g. stability, degradation susceptibility^50^, precipitation tendencies^51^) beyond those encoded by mRNA alone, confounding this comparison. However, serum IgG VH levels do reflect *IGHV* codon optimality, and intentional deoptimization of synonymous V(D)J amino acids sequences consistently reduces expression yield. We conclude that antibody genes have evolved to support diversity and affinity maturation by somatic mutation targeting beyond the benefits for secretion that would be provided by codon optimization.

### Limitations of the Study

Although our results are supported by most optimality metrics tested, there is some discrepancy. Specifically, CSC^F^ averages tended to provide disparate results compared to CSC^W^ & CSC^N^ averages. However, the latter are based on the means of six^16^ and eight^17^ human CSC datasets, respectively, and show strong agreement with each other, while the former is based on one assay^18^ and may be more affected by experimental variability. In addition, CSC-based metrics sometimes provided opposing results to CUF-based metrics (e.g. *IGK* optimality/mutation correlations). While this precludes unambiguous optimality statements in such contexts, these represent two distinct views on optimality (contribution to mRNA stability *vs.* evolved usage frequency) and accordingly, provide complementary insights.

## Methods

### Germline Datasets

Germline *IG* and *TR* reference sequences were downloaded from the International Immunogenetics Information System^1^ (IMGT; imgt.org/vquest/refseqh.html#VQUEST). For regional analyses, available *IGHV*, *IGKV* & *IGLV* FRs/CDRs, as well as *IGHV* “L-PART1+L-PART2” spliced leader and “V-EXON” sequences were downloaded individually. V gene sequences (*IGHV*, *IGKV*, *IGLV*, *IGIV*, *TRAV*, *TRBV*, *TRGV*, and *TRDV*, including leader sequences) were retrieved as “L-PART1+V-EXON artificially spliced sets” (leader sequence spliced with V exon), human C gene sequences (*IGHC*, *IGKC*, and *IGLC*) as “Constant gene artificially spliced exons sets”, and the remainder (*IGHD*, *IGHJ*, *IGKJ*, *IGLJ*, *IGIJ*, *IGIC*, *TRAJ*, *TRBD*, *TRBJ*, *TRGJ*, *TRDD*, and *TRDJ*) as “V-REGION, D-REGION, J-REGION, C-GENE exon sets”. Only “functional”, 5’ complete sequences were analyzed (with the exception of pseudogene/ORF comparisons); 3’ partial sequences were maintained if partiality was <3 codons. Functional V segments that did not begin with ATG start codons were omitted. For D segments, +1, +2, and +3 ORFs were generated; all productive frames were considered to account for frame variability induced by junctional diversification. For V, J, and C segments, only frames corresponding to conserved amino acid sequences were considered.

### Experimental BCR Datasets

Five natural BCR datasets were analyzed; human tonsil, MLN and mouse splenocyte data was generated in-house, while human BM/ALN^45^ and whole zebrafish^48^ datasets were public. Left and right tonsils from a 16 year-old female tonsillectomy patient were collected at NewYork-Presbyterian Hospital, with consent and institutional review board (IRB) approval. After collection, tonsils were combined, minced, dissociated through 100µm strainers to form single cell suspensions, and frozen in 10% DMSO/90% fetal bovine serum (FBS). Mediastinal lymph nodes (MLNs) were collected from a 54 year-old male transplant-inviable organ donor at the University of Chicago with IRB approval. Stations 2L, 4R, 7, 10L, 10R, 12L, and 12R were dissociated as above and frozen separately. For murine splenocytes, five mice were infected with 10 PFU of influenza A/Netherlands/602/2009 H1N1 with IUCAC approval. After 34 weeks, spleens were collected, pooled, dissociated, and stored as above. After thawing, human cells were blocked with Human TruStain FcX Fc Receptor Blocking Solution (Biolegend, #422302, 1:100) in FACS buffer and stained with anti-hCD19-PE-Cy7 and anti-hCD138-PE-Cy7 (Biolegend, #302216 and #356513, 1:100). After washing, cells were incubated with DAPI (Millipore Sigma, D9542, 1:500). Total B cells were identified as DAPI^−^, CD19^+^ and/or CD138^+^ and sorted using a BD FACS Melody. For mouse cells, cells were blocked with anti-mCD16/32 (Biolegend #101302, 1:100) before staining with anti-mCD3-PB and anti-m/hB220-AF488 (Biolegend, #100214 and #103225, 1:100). After washing, cells were incubated with 7-AAD (Biolegend #420404, 1:20). Total B cells were identified as DAPI^−^CD3^−^B220^+^ and sorted using a BD FACS Melody. Sorted cells were loaded onto a 10x Genomics Chip N (#1000375) and emulsions were generated using a Chromium X controller. 5’ gene expression and V(D)J libraries were constructed as per manufacturer instructions (CG000424-RevC) using either human or murine V(D)J enrichment kits, and sequenced using a NextSeq1000 P2 flow cell (Illumina). Single cell RNA sequencing data was processed as described previously^63^, with the following modifications: CellRanger v7.1.0 was used for fastq file processing; GRCh38-2020-A and cellranger-vdj-GRCh38-alts-ensembl-7.1.0 reference genomes were used to align human transcriptomic and V(D)J reads, respectively; refdata-gex-mm10-2020-A and refdata-cellranger-vdj-GRCm38-alts-ensembl-7.0.0 reference genomes were used to align mouse transcriptomic and V(D)J reads, respectively; and Seurat v4.3.0 was used for data analysis. MLN stations were pooled into a single dataset.

In terms of public data, BM/ALN reads (Kim *et al.*^45^, each multiple donors) was downloaded from Zenodo (DOI:10.5281/zenodo.5895181), while whole zebrafish data (Weinstein *et al*.^48^; single whole fish) was acquired from the NIH Sequence Read Archive (Accession SRX003632).

### V(D)J Parsing and Pre-Filtering of Experimental Datasets

For human experimental datasets, V(D)J parsing was performed using VGenes, an in-house BCR analysis program (wilsonimmunologylab.github.io/VGenes). VGenes integrates pre-compiled IgBlast (NIH; ncbi.nlm.nih.gov/igblast) to analyze V(D)J-containing BCR sequences and compute numerous outputs, such as mutational load, isotype, gene usage, sequence quality, and germline reversion. Similarly, pre-compiled RAxML (cme.h-its.org/exelixis/web/software/raxml) was used for phylogenetic analysis of clonal families. Complete V(D)J sequences output by VGenes range from FR1 to FR4, without leader sequences, but with residual C region 5’ fragments. To remedy this, sequences were trimmed at the 3’ end to the nucleotide indicated by IgBlast as the end of the FR4/J region, resulting in strict V(D)Js. Subsequently, sequences were filtered for quality; any that were out of frame, non-productive, negative strand, or possessed a premature stop codon according to IgBlast were filtered out. In addition, any sequences that were incomplete (containing internal “–“ values) or out of frame due to insertions/loose nucleotides not detected by IgBlast (those at sequence start, just prior to FR1) were also omitted. In total, all filtered sequences represented a minor fraction of each dataset.

Due to the inability of IgBlast to map zebrafish V(D)J reads, experimental data from *Danio rerio* were parsed using IMGT/HighV-QUEST (imgt.org/HighV-QUEST/home). *IGH* reads from Weinstein *et al*.^48^ were partial, representing incomplete VDJs. However, many sequences contained complete FR3 and CDR3 sequences. Thus, we analyzed CDR3 CSCs with respect to mutational load within FR3s (CDR3s cannot be germline reverted due to junctions, while FR3s can). Thus, sequences were refined down to those that were productive and possessed intact FR3s & CDR3s.

### CSC averages, CUF averages, CAIs

For humans, individual codon stability coefficient (CSC) values used here were calculated by taking an average of the six sets of experimental CSC values determined across HEK293T, HeLa, RPE, and K562 cells by Wu *et al*. (original publication Source Data 2)^16^, the average of eight sets of CSC values determined in HEK293T cells by Narula *et al*. (original publication Table S5)^17^, or the single set of CSC values determined in HeLa cells by Forrest *et al.* (original publication Table S2)^18^. CSC^Wu^ averages were selected as the primary human readout as they were supported by the most rounded aggregate dataset^16^, spanning six results per codon across three assay types and four human cell types. For mice, individual codon CSC values were calculated using experimental CSC values determined in murine epiblast cells by Forrest *et al.* (S2 Table)^18^. For zebrafish, CSC values were extracted from Figures 1D/F in Bazzini *et al.*^20^ using PlotDigitizer. An average was taken from values extracted from the two graphs to generate the final CSC values. Codon usage frequency (CUF; human & chicken) values were downloaded as codon frequencies per 1000 codons among all RefSeq data from the FDA-associated CoCoPUTs database^27^.

For CSC & CUF averages, means were calculated by breaking ORFs into individual codons, assigning each codon its value, and taking an average over the length of the sequence. Arithmetic means were used given the presence of negative CSC values (incompatible with geometric means).

Codon adaptation indices (CAIs) were calculated as per Sharp & Li^26^. For each codon, CoCoPUTs codon usage frequencies were used to calculate relative synonymous codon usage (RSCU) values (frequency of use / expected frequency of use [1/# of synonymous codons per given amino acid]). Then, RSCU^Max^ values were determined for each codon (the maximum possible RSCU among synonymous codons). For each sequence (germline or natural), the geometric mean of RSCU values for all codons was calculated, along with the geometric mean of RSCU^Max^ values for the same sequence. In both cases, ATG and TGG were excluded from calculations given their role as the sole codon for their respective amino acids, and resulting RSCU/RSCU^Max^ values of 1. Finally, CAIs were calculated as the ratio of (GeoMean RSCU / GeoMean RSCU^Max^).

### Proteomic identification of serum IgG antibody repertoire in flu vaccinees

*BCR-Seq:* N=15 middle-aged female healthcare subjects were selected from a larger clinical cohort reported previously^64^ and received quadrivalent seasonal influenza vaccinations (Fluzone, Flucelvax, or Flublok) during the 2018-2019 season. VH-only and paired VH:VL amplicons were generated using RNA extracted from ≥ 10^6^ PBMCs collected on day 7 post-vaccination, a timepoint at which peak plasmablasts are enriched with high frequency following viral infection or immunization^65^. Bulk VH transcripts were amplified using (RT)-PCR and VH-specific primers, as previously described^31,66^. In parallel, VH:VL paired amplicons were generated from ≥ 2×10^6^ PBMCs that were single-cell isolated in water-in-oil emulsions by a custom flow-focusing device, followed by amplification via overlap extension RT-PCR and nested PCRs using multiplexed VH and VL specific primers, as previously described^67^. The VH and paired VH:VL amplicons were sequenced by Illumina MiSeq 2×300 bp with a minimum 1×10^6^ of reads by the Genomic Sequencing and Analysis Facility at UT Austin Center. After V, D, and J gene annotation using MiXCR^68^, clonotypic lineage was defined by clustering VH sequences based on >90% CDRH3 amino acid identity, measured by Levenshtein distance, using a single linkage hierarchical clustering method as previously reported^31,33^. VH reads sequenced by VH-only or VH:VL paired BCR-Seq were combined and used to create a custom donor-specific MS search database to identify serum IgG antibodies via proteomics Ig-Seq.

*Ig-Seq:* The clonotypic identity and respective abundance of individual serum IgG antibodies were determined by Ig-Seq proteomics, as reported previously^31,33,66,69^. IgG antibodies were purified from 1ml of sera before (day0) and after (day 28) vaccination and then applied to affinity chromatography with immobilized A/Singapore/INFIMH-15-0019/2016 HA trimer antigens (H3 vaccine strain; Native Antigen Co), as previously described^31,33,66^. Briefly, antigen-specific and non-specific antibody fractions were purified from the eluate and flow-through of the antigen affinity column. ELISA binding assay coated with the HA antigen further confirmed the enrichment of antigen-binding fractions in the elution and depletion in the flow-through. For bottom-up LC-MS/MS peptide identification, the eluate and flow-through samples were denatured, reduced, alkylated, and digested into tryptic peptides. The resulting peptides were then injected into a Thermo Ultimate 3000 RSLCnano UPLC coupled to a Thermo Fisher Scientific Orbitrap Fusion Tribrid mass spectrometer by the UT Austin Center for Biomedical Research Support Biological Mass Spectrometry.

CDRH3 spectratyping was conducted, as previously described^31,33,66^. In brief, CDRH3 peptides derived from serum IgG antibodies were identified using the donor-specific custom MS search database comprising VH and VH:VL paired reads sequenced by BCR-Seq, as described above. The high-confidence, unique CDRH3 peptides mapped to a single clonotypic lineage were used to quantify the abundance of individual antibody clonotypes by summing the extracted ion chromatogram (XIC) peak area of associated CDRH3 peptides. The % VH gene abundance was determined by summing the abundance of individual clonotypes using the same VH gene and averaged across all fifteen vaccinees. These values, calculated separately from HA-specific or non-specific fractions, were then correlated against CSC values for germline *IGHVs*.

### Mutability Scoring

Mutability score averages (MutScores) were generated using the SHazaM package^39^ in R (shazam.readthedocs.io). Briefly, sequences were broken down into iterative 5-mers, and a MutScore was estimated for the central nucleotide based on empirically-derived human heavy chain data. The scores for each nucleotide were then averaged over the length of the sequence. The first and last two nucleotides of each sequence were not included in this calculation given that they were missing 1-2 nucleotide neighbours each (upstream and downstream, respectively) and could not comprise the central position in a 5-mer.

### *IGHV* Mutability Modification

To generate maximally-mutable variants, six germline *IGHV* sequences (*IGHV4-28*01*, *IGHV5-51*01*, *IGHV4-38-2*01*, *IGHV3-49*01*, *IGHV3-64*01*, *IGHV3-15*01*) were selected. After separating sequences into FRs & CDRs, each region was converted to its corresponding amino acid sequence using Python. Synonymous codons were considered for each residue and all possible permutations which maintained the original amino acid sequence were generated. The resulting set of candidate sequences was analyzed for predicted mutability using the SHazaM package. After selecting the regions with the highest averaged MutScores, FRs, CDRs, and leader sequences were rejoined, and CSC averages were calculated for final *IGHV* sequences. This process was performed in two iterations for each gene, one where only CDRs (MaxMut^CDR^) were modified, and one where both FRs & CDRs (MaxMut^FR/CDR^) were modified. To generate *IGHV*s with standard human codon usage frequencies, amino acid sequences were imported into SnapGene and reverse translated using human codon tables. Sequences for modified IGHVs are presented in **Table S4**.

### SHM Simulation

To predict the effect of SHM on codon optimality, we ran immuneSIM^44^ (immunesim.readthedocs.io) on *IGH*, *IGK* and *IGL* human reference repertoires using the parameters outlined in **Table S5**. Eight iterations were run for each chain with varying SHM probability, after which datasets pooled per chain in order to get a maximal range of mutations. V(D)J information and mutational load were parsed using VGenes and filtered as described above for natural datasets.

### Constant Region Buffering Analysis

To assess whether constant regions buffer V(D)J codon optimality, CSC averages were calculated for *IGH*, *IGK* and *IGL* sequences from tonsil B cells in isolation (VDJ^FR^^1-FR4^) and after splicing with V-matched leader sequences, with or without constant region exons (IMGT).

### mAb Expression Testing

Heavy and light chain plasmid vectors were generated as described previously^70,71^; briefly, V(D)J^FR1-FR4^ sequences derived from MLNs (set 1) or PBMCs (set 2; derived from a previously-reported dataset^71,72^ archived under GEO accession numbers GSE171703 and GSM5231088–GSM5231123) were spliced with upstream/downstream Gibson cloning fragments, ordered from Integrated DNA Technologies (IDT), and cloned into AbVec hIgG1, hIgK or hIgL expression vectors. For standard mAb expression studies, plasmids were transiently transfected into Expi293F cells (5ml cultures in 50ml bioreactors [Corning Mini Bioreactor Centrifuge Tubes, FisherScientific #07-202-150], 3.0e6 cells/ml) at a ratio of 1:1 heavy/light chain (5µg+5µg) in OptiMEM+Expi293Reagent using manufacturer’s protocol. Cells were then incubated for five days at 175rpm, 37°C with 8% CO_2_. To assess IgG expression, cultures were centrifuged at 3000rcf for 10min/4°C to pellet cells. Supernatants were decanted, frozen at –20°C, and subsequently analyzed for total hIgG by ELISA. Briefly, plates (Corning 96 Well EIA/RIA Assay Microplate, ThermoFisher #3369) were coated with goat anti-hIgG(H+L) antibody (ThermoFisher #62-8400, 1µg/ml in PBS) overnight at 4°C. The next day, coating antibody was removed by flicking and plates were blocked with 200µl 1% BSA in PBS for 1h at room temperature. After washing six times, plates were incubated with 50µl supernatant diluted 1:5000 or 1:100 (as necessary for assay linear phase) in 1% BSA. hIgG1 purified in-house was used as a standard. After 1h, plates were washed six times, and incubated with 50µl of anti-hIgG(Fc)-HRP detection antibody (Jackson Immunoresearch #109-035-098, 1:1000). After a further 1h, plates were washed and developed with 50µl SuperAqua Blue solution (Invitrogen #00-4203-58). Plates were read at 405nm after 2-3min.

For pDNA transfection mass titration work, heavy and light chain pDNA was combined at 5µg+5µg, serially diluted in OptiMEM as indicated in **Figure 5f**, and transfected with a fixed amount of Expi293Reagent. For deoptimized mAb titration studies, transfection was modified to include differing ratios of original and deoptimized heavy or light chain plasmids as indicated in **Figure 5g**. Expression was then analyzed by ELISA as described above.

## Statistical Analyses

All statistical analyses were conducted with GraphPad Prism v.10.0.2 with the following exceptions: 1) Multivariate regression was performed using the Scatterplot3d R package; 2) Microsoft Excel was used for bulk, Bonferroni-corrected Pearson correlations between VDJ mutations and CSC averages within spike^+^ clonal families. One-way ANOVA with Tukey or Dunnett’s test was used to analyze one-factor multi-group comparisons; repeated measures ANOVA with Greenhouse-Geisser correction was used for paired instances. (Un)paired t-tests were used to analyze two-group comparisons. One-sample t-tests were used to compare single datasets to a null hypothesis (theoretical mean of 0). Linear regression with Pearson correlation was used to analyze bivariate correlations. Specific tests are indicated in figure legends. Significance (*) was defined as p<0.05.

## Lead contact

Further information, and requests for resources and data, should be directed to and will be fulfilled by the lead contact Dr. Patrick C. Wilson.

## Materials Availability

This study did not generate new unique reagents.

## Data and Code Availability

Raw sequencing data have been deposited to the NCBI Gene Expression Omnibus (GEO) under accession numbers GSE260943 (human tonsil B cells; GEXLIBs/VDJLIBs), GSE260944 (human MLN B cells; GEXLIBs/VDJLIBs), and GSE260942 (mouse splenocyte B cells; VDJLIBs); data will be publicly available at the time of publication. VGenes, and all other original codes used in this study, are freely available from the Wilson Lab GitHub page at: https://github.com/WilsonImmunologyLab (see *VGenes* and *Codon-Optimality*). All other data, or information required to reanalyse the data reported in this paper, are available from the lead contact upon request.

## Acknowledgements

This study was supported by funding from the National Institute of Allergy and Infectious Diseases (NIAID; grants P01AI165077, 5U01AI144616, 1U01AI165452, 1U19AI168632, U01AI153700; contracts 75N93019C00051 and 75N93019R00028; PCW), the Centers for Disease Control & Prevention (CDC; 75D30119C06088; GG) and the Bill & Melinda Gates Foundation (INV-004956; GG). Tonsil collection was supported by funding from the Leukemia & Lymphoma Society (LLS 7027-23, LLS-SCOR 7026-21; GI). JJCM is supported by a Banting Postdoctoral Fellowship (BPF-186528) from the Canadian Institutes of Health Research (CIHR). The authors wish to thank Drs. Stefan Worgall and Robert E. Schwartz (WCM) for advice on MLN selection.

## Author Contributions

JJCM and PCW conceived the project and designed the experiments. JJCM, JP, SC, GDW and MH performed experiments. YF, NYZ, JS, and SAN provided support for experiments. JRK and MLLM provided human MLN samples. GI provided human tonsil samples. CAT and JCC performed bioinformatic analyses, SHM simulations, V(D)J (de)optimizations, and optimality/mutability score calculations. LL and PCW developed VGenes. JJCM and JP performed data analysis. JJCM, PCW and GG supervised the study and interpreted the data. JJCM drafted the manuscript. All authors provided feedback and approved the final version of the manuscript. PCW and GG acquired funding.

## Supplemental Information

Supplemental information is available online.

- **Figure S1**: Extended data for Figure 1.
- **Figure S2**: Extended data for Figure 2.
- **Figure S3**: Extended data for Figure 3.
- **Figure S4**: Extended data for Figure 4.
- **Figure S5**: Extended data for Figure 5.
- **Figure S6**: Annotated correlation between Wu *et al.* mean CSC values and CoCoPUTs CUFs.
- **File S1**. Supplementary tables:

○ **Table S1**. Values used to calculate CSC averages, CUF averages, and CAIs.
○ **Table S2.** Compiled sequences for highly-expressed proteins and multigene families.
○ **Table S3**. Codon optimality rankings for individual human IGHV alleles as determined by CSC averages, CUF averages, and CAIs.
○ **Table S4**. Original and mutability-modified IGHV sequences.
○ **Table S5**. immuneSIM analysis parameters.
○ **Table S6**. Codon enrichment/depletion in highly-mutated sequences.
○ **Table S7**. mAb V(D)J sequences.

---

**Figure S1. Extended data for Figure 1.**
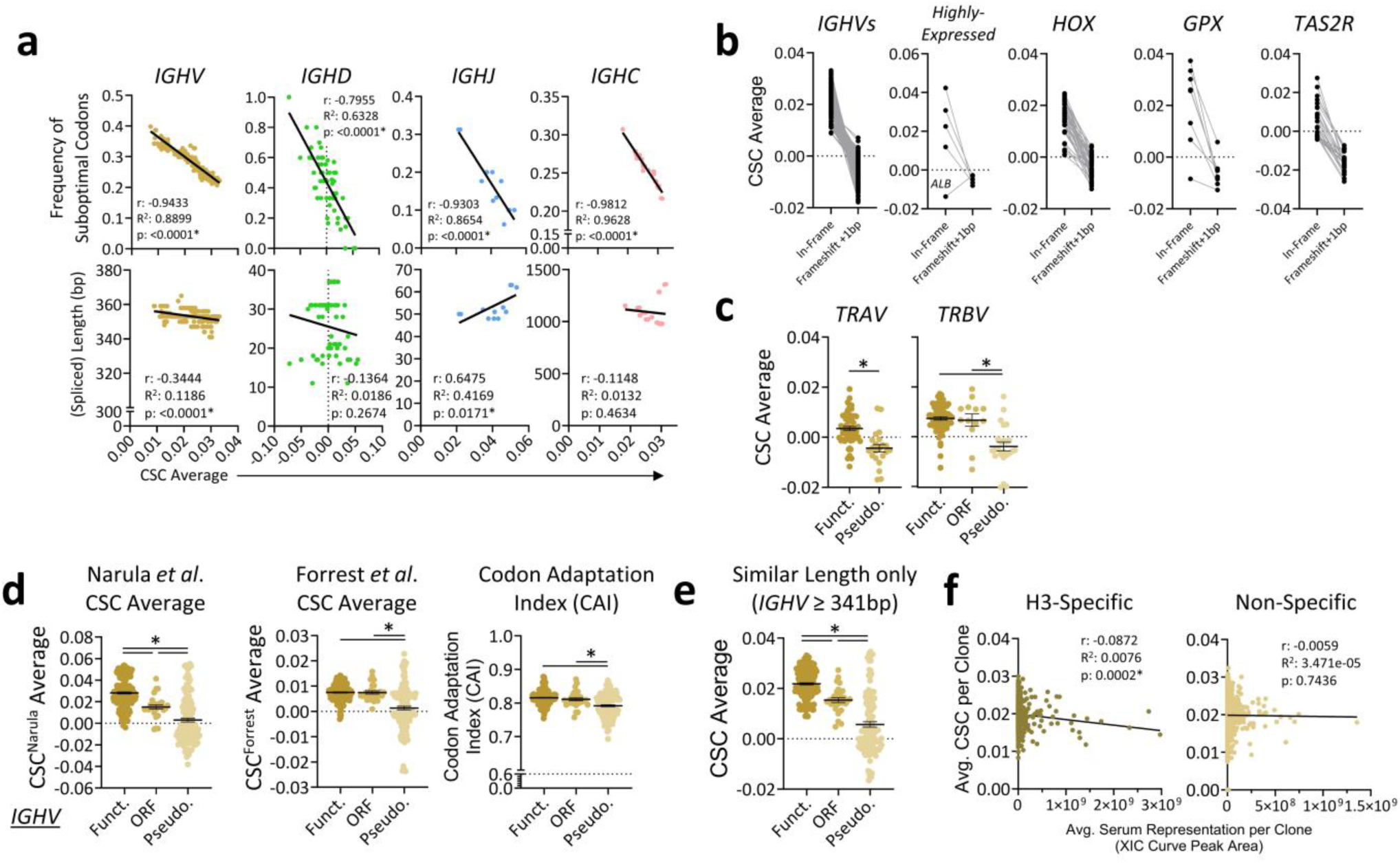
(**a**) Pearson correlations between CSC averages and (top) frequency of suboptimal codons or (bottom) spliced length for *IGHV*, *IGHD*, *IGHJ* and *IGHC* alleles. **(b)** Paired CSC averages for original and +1bp frameshifted *IGHVs* and control coding sequences. **(c)** CSC averages for functional, ORF-designated and pseudogene *TR* V gene variants. **(d)** CSC^N^ averages, CSC^F^ averages, and CAIs were compared between functional, ORF-designated and pseudogene IG V gene variants. **(e)** CSC averages for functional, ORF-designated and pseudogene *IGHV* variants of similar length were compared. **(f)** Pearson correlations between mean CSC average vs. average IgG clonal serum representation, for matched clones, separated into H3^+^ and H3^−^ fractions. **Data: (c,d,e)** mean ± SEM. **Statistics: (a,e)** linear regression with Pearson’s correlation; **(c,d,e)** one-way ANOVA with Tukey’s test. *: p<0.05.

**Figure S2. Extended data for Figure 2.**
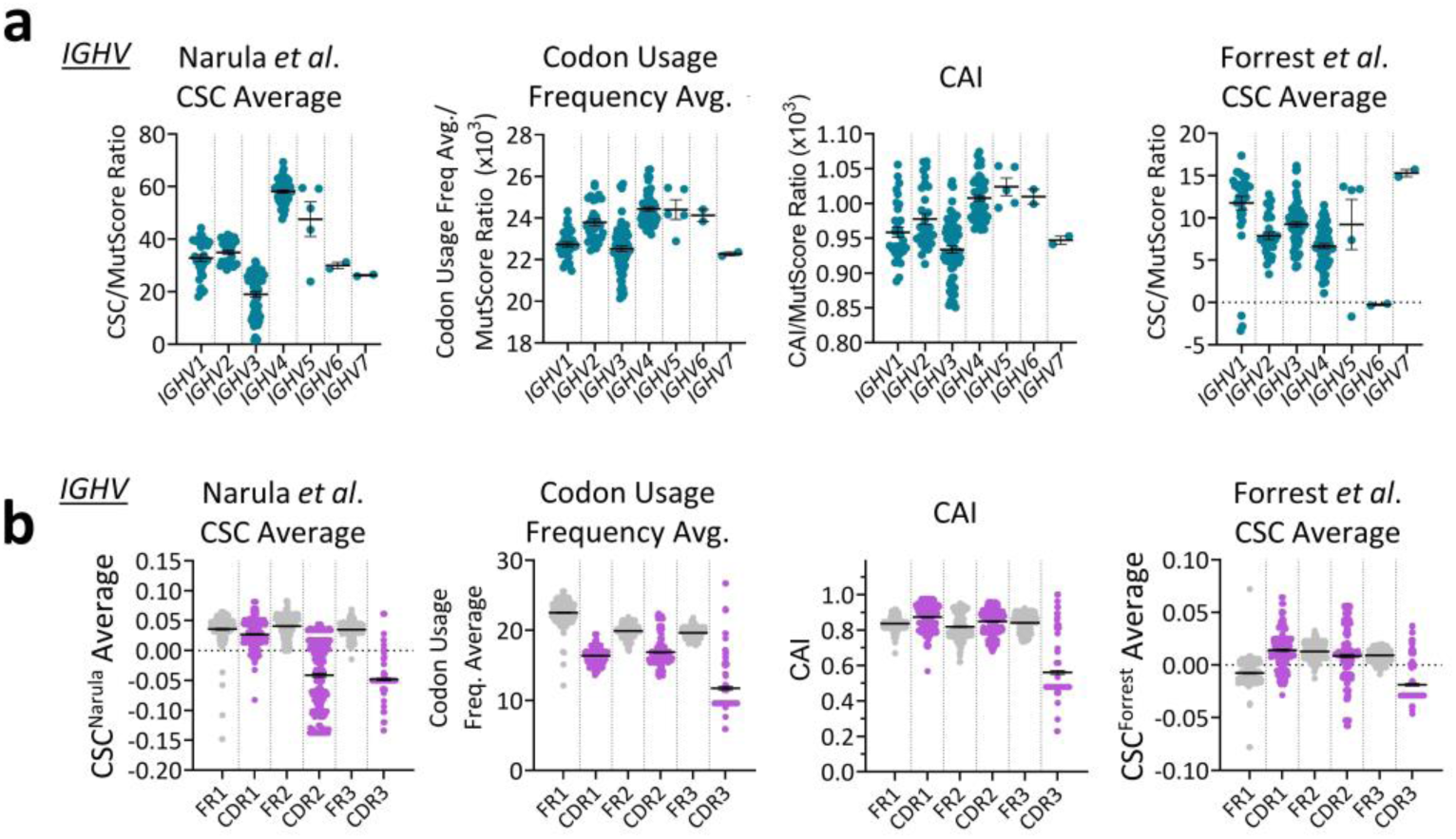
(**a**) Optimality/MutScore ratios were calculated for each *IGHV* gene family in the form of CSC^N^ averages, CSC^F^ averages, CUF averages, and CAIs. **(b)** Available isolated FR and CDR regions were downloaded for each *IGHV* gene sequence, and optimality scores were calculated in the form of CSC^N^ averages, CSC^F^ averages, CUF averages, and CAIs. **Data:** mean ± SEM.

**Figure S3. Extended data for Figure 3.**
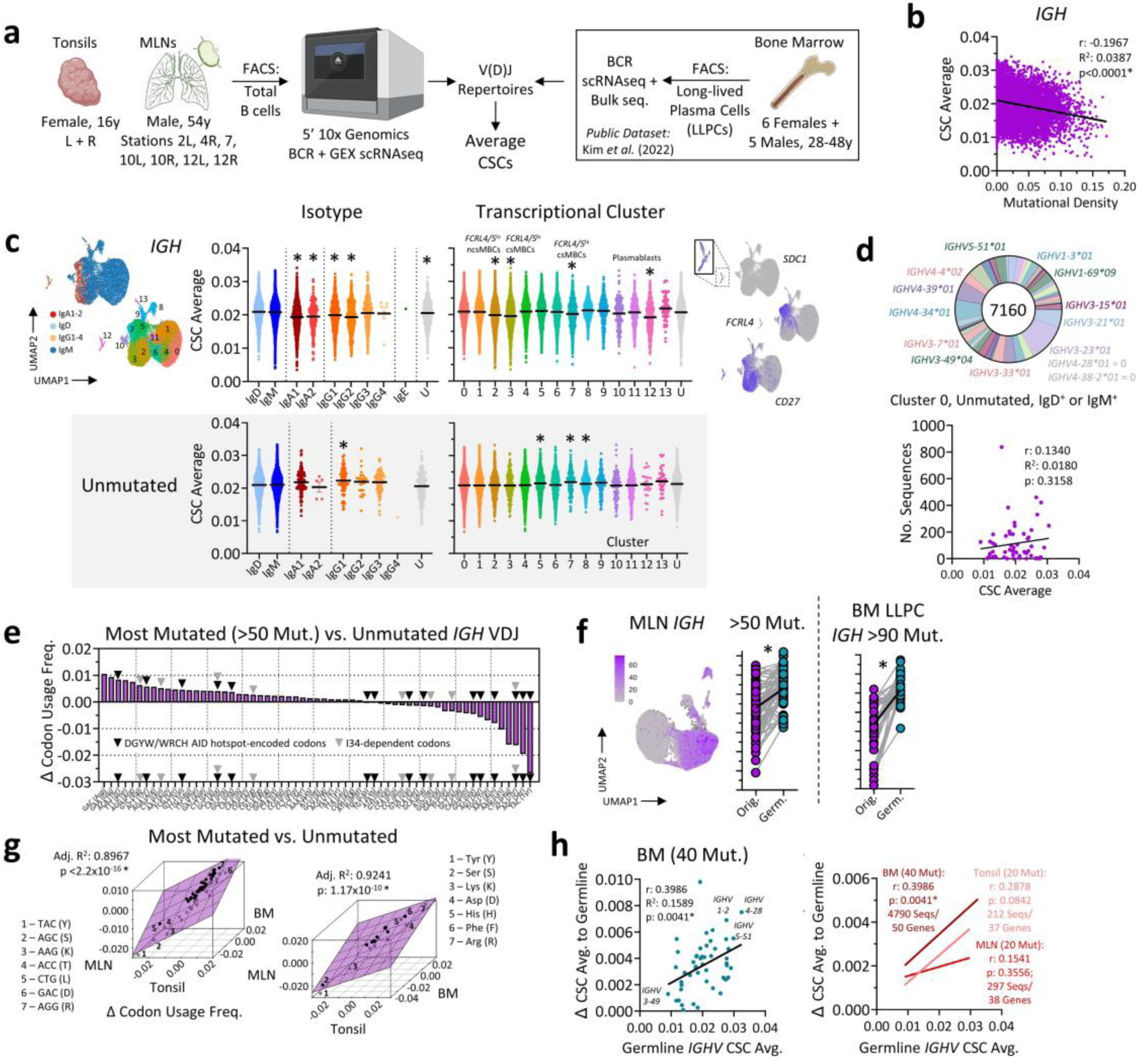
(**a**) Schematic demonstrating the overall workflow for analysis of natural V(D)J repertoires from human tonsil B cells, mediastinal lymph node (MLN) B cells, and bone marrow plasma cells. **(b)** Tonsil *IGH* V(D)J CSC averages were correlated against mutational density (load/bp). **(c)** UMAP at left demonstrates isotype and transcriptional subclusters within tonsil B cells. *IGH* V(D)J CSC averages were determined, and broken down by isotype (left) or transcriptional subcluster (right). Specific gene feature plots are shown at far right to landmark select cell types of interest; *SDC1* identifies plasmablasts/plasma cells, *CD27* identifies memory B cells, and *FCRL4* differentiates specific memory subsets. This analysis was conducted on (top) all V(D)Js or (bottom) unmutated V(D)Js only. **(d)** Left: Breakdown of cluster 0, unmutated, IgD^+^ or IgM^+^ cells in terms of *IGHV* gene usage. Right: Pearson correlation between number of sequences used per *IGHV*, with *IGHV* CSC average. **(e)** Tonsil *IGH* sequences with >50 or zero mutations were pooled separately; codons were counted for each pooled group, and usage frequencies were calculated. Change (Δ) in usage is plotted. **(f)** Matched original and germline CSC averages were plotted as paired data for highly-mutated MLN and BM IGH sequences. UMAP at left shows *IGH* mutation distribution among MLN B cells. **(g)** Multivariate regression demonstrating change in (left) codon usage or (right) amino acid usage between highly-mutated and naïve tonsil, MLN, and BM *IGH* sequences. **(h)** Left: BM plasma cell *IGH* sequences were filtered to only include those with exactly 40 mutations (3986 sequences). Subsequently, the change in CSC average between mutated and germline variants of each sequence was calculated, pooled based on *IGHV* gene, and a mean was calculated per *IGHV* gene. This mean was then correlated against the germline CSC averages for individual *IGHV* genes (mean of alleles). Right: The same analysis was conducted in tonsil cells (filtered to exactly 20 mutations) and MLN cells (filtered to 20 mutations). **Statistics: (b,d[right],h)** linear regression with Pearson’s correlation; **(c)** One-way ANOVA with Tukey’s test, significance relative to IgD; **(f)** Paired t-test. **(g)** Multivariate regression. *: p<0.05. Graphics in **(a)** from Biorender.com.

**Figure S4. Extended data for Figure 4.**
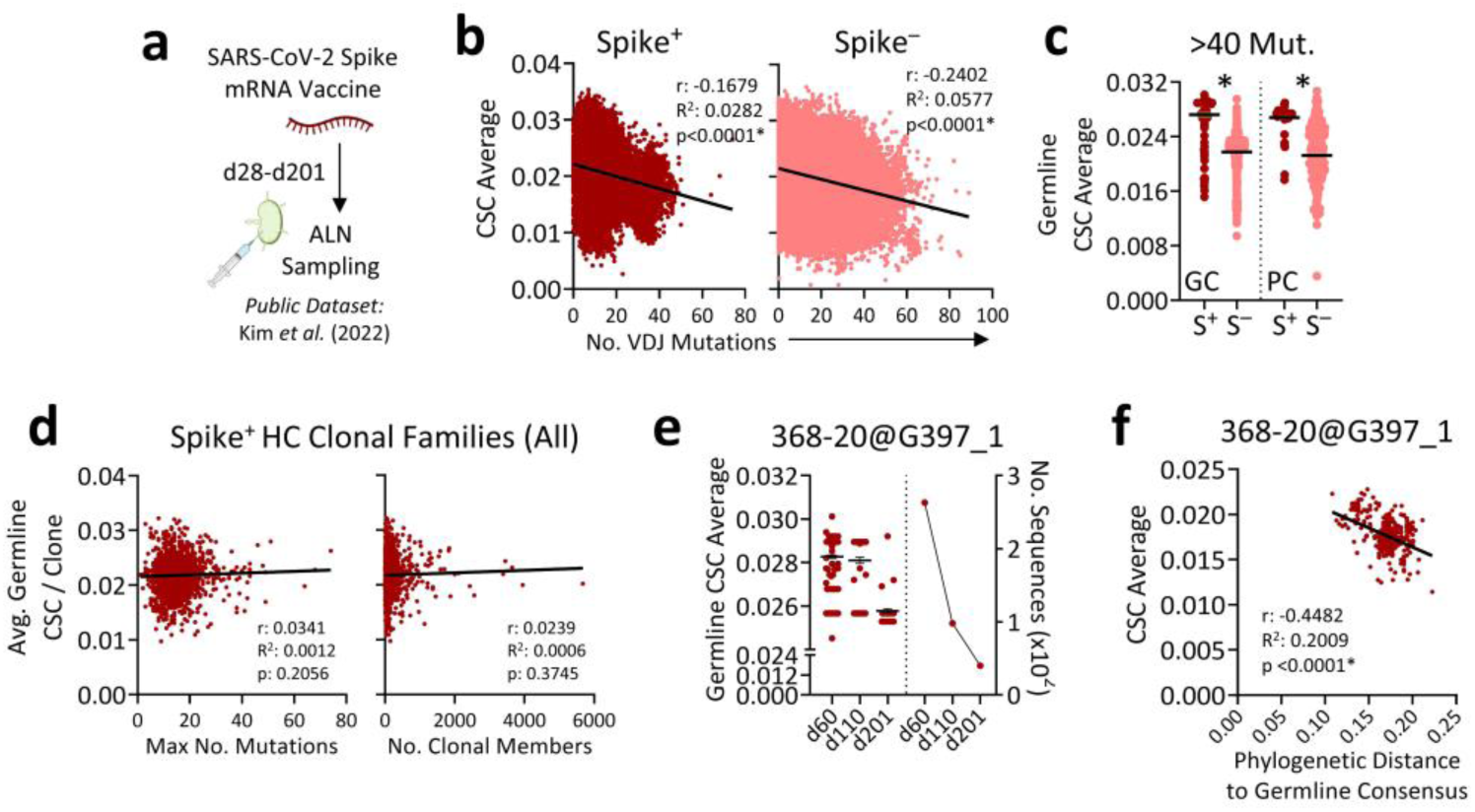
(**a**) Schematic demonstrating the overall workflow for analysis of natural VDJ repertoires from human axillary lymph node (ALN) B cells following SARS-CoV-2 mRNA vaccination, as performed by Kim *et al*. (2022). **(b)** Pearson correlation between CSC averages and the number of mutations per IGH sequence, separated into SARS-CoV-2 spike^+^ and spike^−^ fractions as per the above publication^45^. **(c)** Germline CSC averages were plotted for spike^+^ and spike^−^ *IGH* sequences of germinal center (GC) B cells and plasma cells (PCs) with >40 mutations. **(d)** The average germline CSC average per spike^+^ heavy chain clonal family was correlated against (left) the maximum number of mutations or (right) number of clonal members per *IGH* clonal family. **(e)** Left: germline CSC averages were plotted for members of the *IGH* clonal family 368-20@G397_1^45^, broken down by day post-vaccination. Right: the number of sequences belonging to this family is plotted over time. **(f)** For each member of the *IGH* clonal family 368-20@G397_1^45^, CSC averages were correlated against phylogenetic distance to germline consensus. **Data: (c,e[left])** mean ± SEM. **Statistics: (b,d,f)** linear regression with Pearson’s correlation; **(c)** unpaired t-tests. *: p<0.05. Graphics in **(a)** from Biorender.com.

**Figure S5. Extended data for Figure 5.**
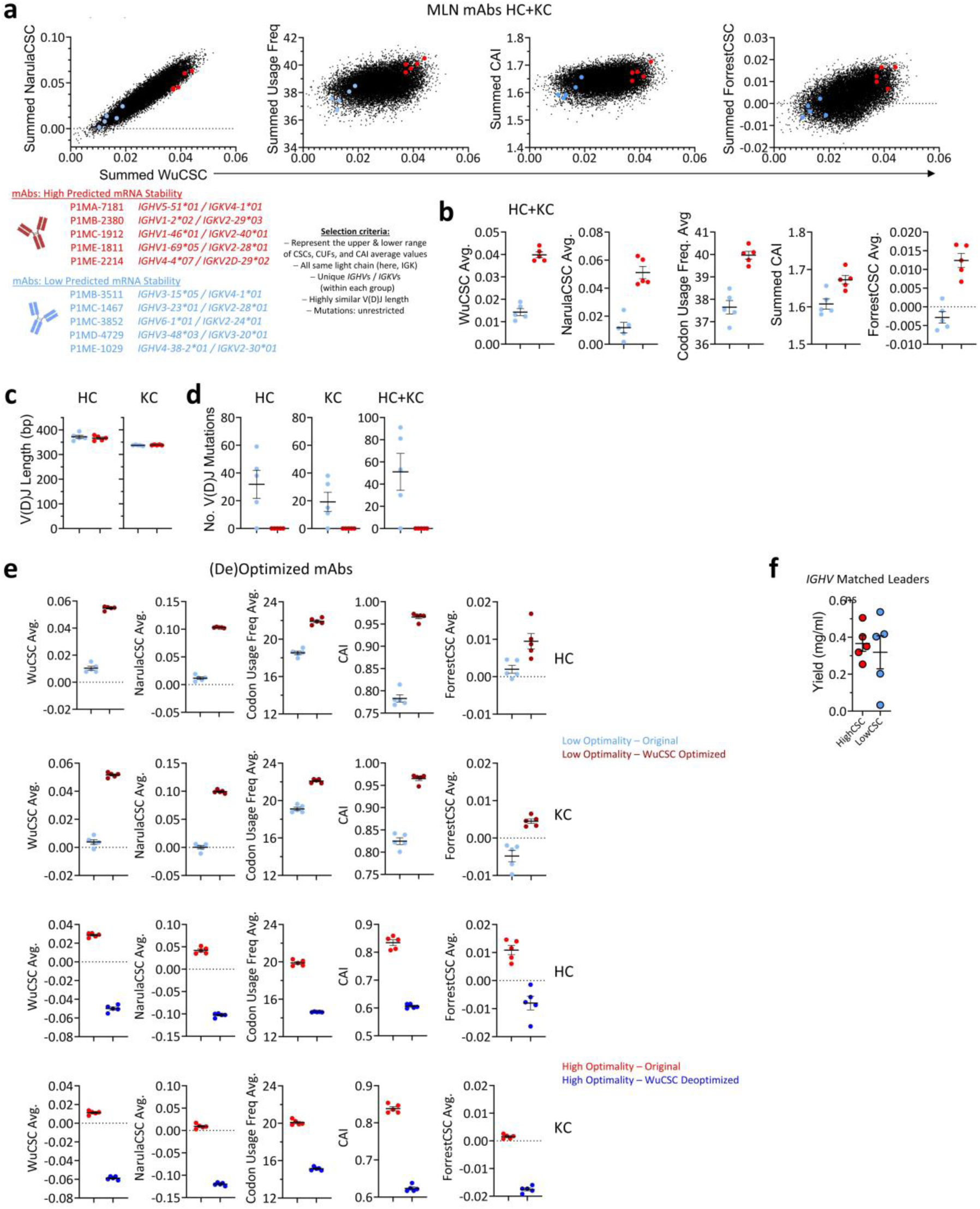
Set 1 mAb selection and characterization. **(a)** Summed CSC^W^ averages for heavy and light chains were calculated for MLN B cell clones, and correlated against summed CUF averages, CAIs, CSC^N^ averages, CSC^F^ averages. Five clones with high (red) and five with low (blue) optimality scores were selected, with further criteria as detailed in black text. **(b)** Summed optimality scores for selected clones. **(c)** V(D)J lengths for selected clones, by chain. **(d)** Number of V(D)J mutations in the selected clones, by chain and combined. **(e)** Optimality scores for optimized and deoptimized variants of the selected clones as compared to original, separated by chain. **(f)** Alternate versions of set 1 high/low optimality clones with *IGHV*-matched leader sequences were expressed in Expi293F cells; yield in culture supernatants was quantified by total IgG ELISA. **Data: (b-f)** mean ± SEM. **Statistics: (f)** unpaired t-test. *: p<0.05.

**Figure S6.**
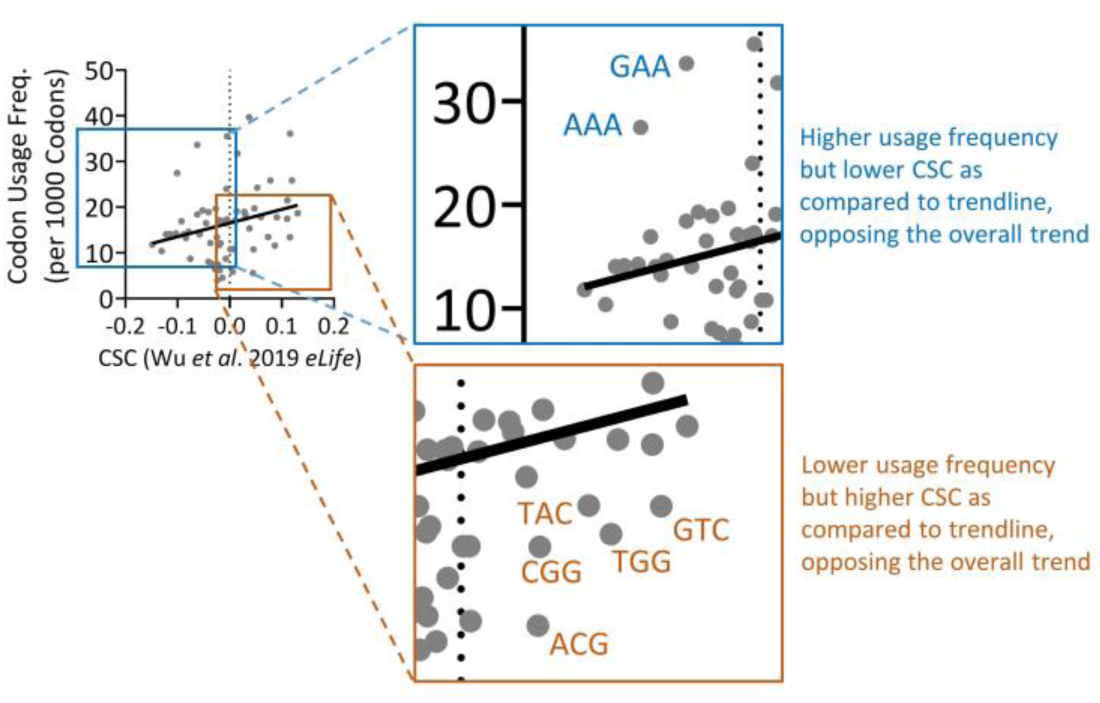
Annotated correlation between Wu et al. mean CSC values and CoCoPUTs CUFs. Pearson correlation between CSC^W^ and CUF base values. Although these datasets correlate positively overall, some codons (highlighted at right) display features in opposition to the overall trend (high CSC but low CUF, or vice versa).

